# Personalized Gut–Liver Microphysiological System Maps Donor-Specific Tissue-Resident Immunity and Reveals a Conserved Metabolic Crosstalk

**DOI:** 10.1101/2025.05.05.652244

**Authors:** Merve Uslu, Ran Ran, Mohd Farhan Siddiqui, Jing Liang, Luther Raechal, Matjaz Dogsa, Carole Perrot, Linda Lieberman, Douglas K. Brubaker, Martin Trapecar

## Abstract

Tissue-resident immune (TRI) niches are unique to tissues and greatly vary between individuals. We built a personalized gut–liver microphysiological system (MPS) to recapitulate these profiles, combining primary colon epithelium, hepatocytes, and autologous CD45⁺ TRI cells of two donors. Single-cell RNA-seq of colon and liver revealed distinct TRI profiles and predicted responses distinct between donors. Co-culture established organ and donor-specific immune programs: colonic epithelium induced Th1/Th17 polarization in Donor 1 but B cell differentiation in Donor 2. Gut-liver crosstalk in all donors converged on a retinoid–bile acid metabolic axis with a muted inflammatory set-point, indicating that circulating metabolites can override baseline immune differences. Microbial agonist challenges of gut compartments revealed distinct liver responses: Poly(I:C) induced a uniform type-I/III interferon burst, LPS triggered a stronger response in Donor 1, and 5-OP-RU selectively activated Donor 2. Our personalized, immune-competent gut–liver MPS demonstrates that a conserved metabolic dialogue coexists with and is modulated by TRI profiles. This work provides a blueprint for exploring immunometabolic diseases and precision therapeutics in multi-organ models reflecting human immune diversity.

## Introduction

The intestine and liver host two critical barrier interfaces with uniquely enriched and complex tissue-resident immune (TRI) populations^1^. Single-cell atlases reveal extraordinary cellular diversity and donor-specificity within these niches^2–5^. TRI cells, including specialized tissue-resident memory (TRM)^6^ cells, Innate Lymphoid Cells (ILCs)^7^, and macrophage subsets^8, 9^, exhibit distinct transcriptional and metabolic programming compared to their circulating counterparts, equipping them for rapid, site-specific surveillance and response^10^. Understanding how this localized, personalized immunity coordinates physiology across organs remains a challenge.

Studying these human systems is hindered by model limitations. Animal models fail to capture key aspects of human-specific immunity and metabolism^11^. 2D cultures lack physiological architecture, but even advanced organoids often omit crucial vascular or mechanical cues and cannot model inter-tissue communication ^12, 13^. Microphysiological systems (MPS) aim to bridge this gap ^14–16^, but many platforms utilize immortalized cell lines or add circulating immune cells, failing to incorporate the complexity and unique conditioning of genuine TRI compartments. Consequently, key determinants shaping tissue immunity such as local antigen experience, stromal education, and metabolic imprinting remain under-represented in vitro^13, 17^. Furthermore, substantial inter-individual immune variation, primarily driven by non-heritable factors^18^, is rarely accounted for.

The gut-liver axis provides a compelling system for investigating such inter-organ dynamics. The portal connection exposes the liver to a concentrated influx of gut-derived nutrients, microbial products (like LPS recognized by TLR4^19^, xenobiotics and bile acids^20^). Perturbations within this circuit are strongly implicated in prevalent pathologies like NAFLD, IBD^21, 22^ and possibly neurodegeneration^23, 24^. Beyond passive transit, bile acids function as signaling molecules via FXR and TGR5, modulating both metabolism and inflammation ^25^. Furthermore, specialized immune cells like Mucosal-associated Invariant T cells (MAIT) cells, abundant in the liver, directly link microbial metabolism via recognition of riboflavin derivatives like 5-OP-RU presented by MR1^26^ to hepatobiliary immunity, exemplifying the axis’s integrated nature^27^.

While initial gut-liver chip models, have demonstrated utility for basic toxicology or metabolism studies^28^, they lacked immunological depth and personalization. These attempts were further improved with models of the gut-liver^29^ and gut-liver-brain^30^ axis where each tissue was accompanied by its macrophage niche and perfused with circulating CD4 T regulatory and Th17 cells. An important recent development was the creation of human intestinal organoids with autologous, tissue-resident immune compartments^31^, albeit limited to the lymphocyte niche. To date, however, no platform has successfully reconstituted autologous gut epithelium, hepatocytes, and their corresponding complex, matched TRI populations from the same donor to interrogate how individual heterogeneity dictates immune crosstalk and functional responses to defined microbial or viral challenges.

Addressing this gap, the present study aimed to develop an autologous, gut-liver-TRI MPS. We integrated primary human cells from paired colon (epithelium, intraepithelial leukocytes and lamina propria TRI) and liver tissues (hepatocytes, liver TRI) from individual donors into the Multiorgan Tissue Interaction Vessel (MOTIVE-2) platform. We combined scRNA-seq atlases with computational ligand interaction modeling^32, 33^ and bespoke epithelial-immune co-cultures based on colonic and liver multilineage TRI, to ask: 1) How do donor-specific TRI landscapes shape local immune tone ex vivo? 2) How are parenchymal cells informing immune signaling hubs? 3) Does fluidic Gut-Liver coupling induce convergent physiological programs? 4) Are early responses to defined microbial/viral stimuli (LPS, Poly I:C, 5-OP-RU) donor- and stimulus-specific, reflecting baseline profiles?

By answering these questions, we provide an approach capable of dissecting both conserved metabolic crosstalk and personalized immune reactions within an autologous human gut-liver immune axis, laying groundwork for precision modeling and therapeutic testing (Fig.1a).

## Results

### 1. Single-Cell Profiling Reveals Diverse Tissue-Resident Immune and Parenchymal Cell Landscapes in Colon and Liver with Stark Differences Across Donors

Prior to TRI-MPS integration, we profiled the tissue-resident immune (Fig. 1) and non-immune (Fig. S1) compartments of a male and female donor by applying single-cell RNA-seq to freshly isolated cells from matched colonic and hepatic tissues. After quality control and integrated clustering of cells, we resolved 26 transcriptionally distinct immune clusters that spanned all major lymphoid and myeloid lineages using Leiden clustering^3, 4^ (Fig. 1b left: colon, right: liver). Marker-gene signatures confirmed canonical identities, including CD4, CD8A, CCR7 and SELL in T-cell subsets, MZB1 and IGKC in plasma cells, and MARCO/VSIG4 in Kupffer-like macrophages, indicating that the workflow captured the breadth of immune diversity present in both organs.

**Figure 1:**
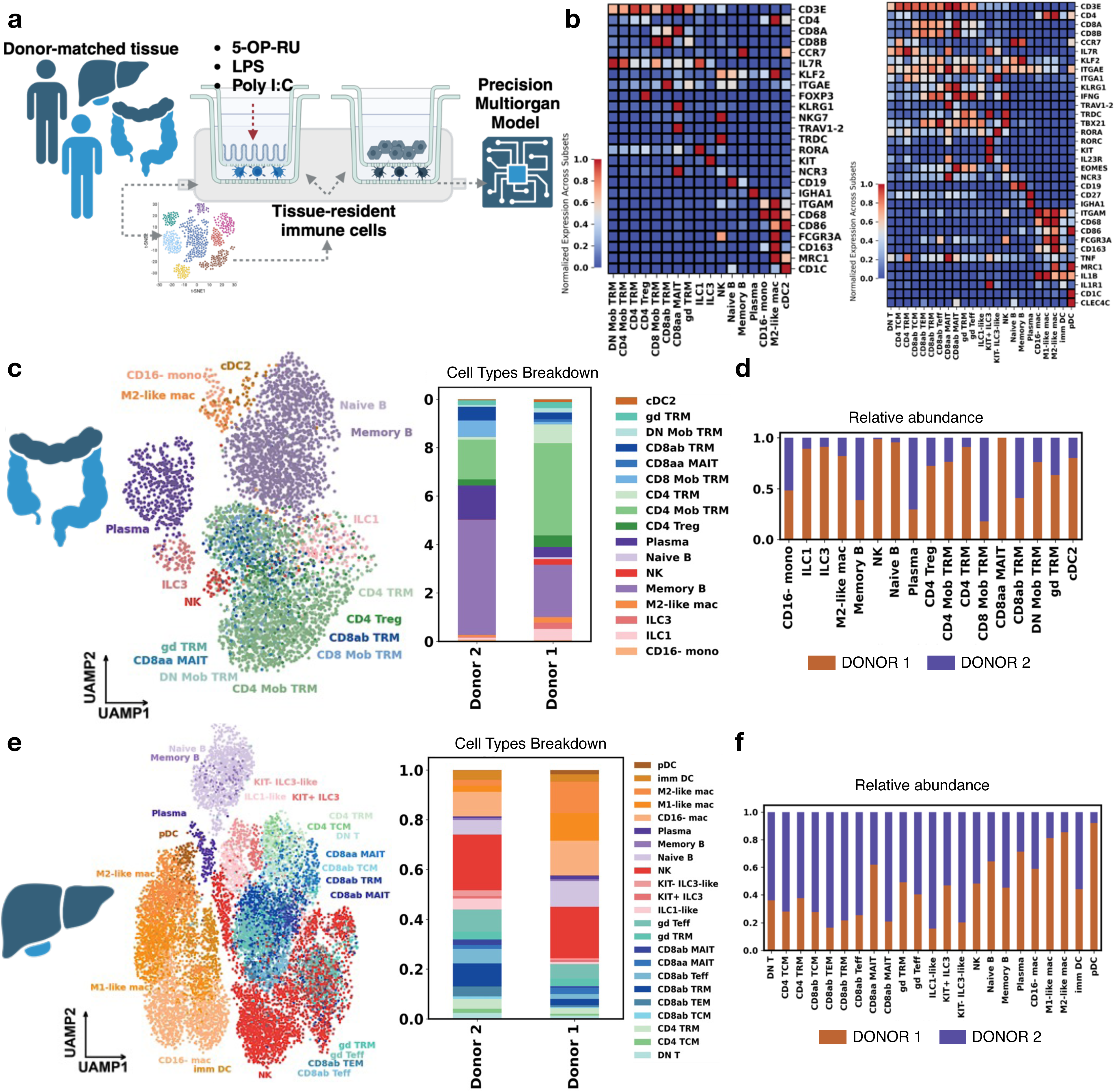
Single-cell characterization of same-donor colon and liver tissue reveals significant differences in the tissue-resident immune repertoire. **a,** graphical representation of the project, approach and its aims. **b,** Marker gene expression in all identified colon (left) and liver (right) immune cell subsets in fresh tissue samples. The color of each cell in the heatmap denotes the normalized cell type-averaged expression of each marker. Abbrevations: Mob - mobile; CD4 TRM - CD4^⁺^ tissue-resident memory T cells; CD4 Treg - CD4^⁺^ regulatory T cells; CD8ab TRM - CD8αβ tissue-resident memory T cells; CD8aa MAIT - CD8αα mucosal-associated invariant T cells; DN T - double-negative T cells; gd TRM - γδ tissue-resident memory T cells; Naive B - naïve B cells; Plasma - plasma cells; Memory B - memory B cells; NK - natural killer cells; ILC1 - innate lymphoid cells type 1; ILC3 - innate lymphoid cells type 3; M2-like mac - M2-like macrophages; CD16-mono - CD16^⁻^ monocytes; cDC2 - Type-2 conventional dendritic cells; CD16-mac - CD16^⁻^ macrophages; imm DC - immature dendritic cells; KIT^⁺^ ILC3 - KIT^⁺^ innate lymphoid cells type 3; KIT^⁻^ ILC3-like - KIT^⁻^ innate lymphoid cells type 3-like cells; M1-like mac - M1-like macrophages; CD4 TCM - CD4^⁺^ central memory T cells; gd Teff - γδ effector T cells; pDC - plasmacytoid dendritic cells; CD8ab TCM - CD8αβ central memory T cells; CD8ab TEM - CD8αβ effector memory T cells; CD8ab Teff - CD8αβ effector T cells; CD8ab MAIT - CD8αβ mucosal-associated invariant T cells. **c,** Left: UMAP embeddings of immune cells collected in the colon of all donors; Right: stacked bar plot shows fractional cell type composition in Donor 1 and Donor 2. **d,** The bar plots show the contribution of each donor to each of the colonic immune cell types in the integrated data and expressed as relative abundance. **e,** Left: UMAP embeddings of cells collected in the liver of all donors; Right: stacked bar plot shows fractional cell type composition in Donor 1 and Donor 2. **f,** The bar plots show the contribution of each donor to each of the hepatic immune cell types in the integrated data and expressed as relative abundance.

Donor identity emerged as the dominant source of variation in the colon. UMAP projection segregated CD4⁺ and CD8⁺ tissue-resident memory T cells (Trm), MAIT and γδ T cells, innate lymphoid cells, plasma cells, B cells and multiple macrophage states (Fig. 1c) in alignment with recent atlases of the human gut^2, 34^. However, their relative abundances differed sharply between donors: Donor 1’s colonic compartment was heavily skewed toward Trm populations and inflammatory macrophages, whereas Donor 2 showed a pronounced expansion of humoral and innate compartments, with plasma cells, MAIT and NK cells making up more than one-third of all leukocytes (Fig. 1d).

A parallel analysis of hepatic CD45⁺ cells reproduced 24 discrete clusters that included Kupffer-like macrophages, CD16⁺ monocytes, pDCs, NK/ILC subsets, B-cell states and multiple effector/memory T-cell subsets (Fig. 1e), mirroring the diversity recently described^3^. Donor-specific biases again predominated: Donor 1 contributed disproportionately to Trm subsets and M1/M2 macrophages, whereas Donor 2 was enriched for plasmacytoid DCs, ILC3-like cells and plasma cells (Fig. 1f). Although the exact lineage skews differed between gut and liver, the data collectively show that inter-individual variation outweighs tissue-of-origin for several key immune lineages.

The paired-tissue pipeline reliably captures the full spectrum of tissue-resident immune niches while revealing highly personalized immune fingerprints that are conserved across organs within each donor. These donor-specific biases in adaptive and innate compartments provide the mechanistic rationale for reconstructing immunocompetent, donor-matched gut–liver models in downstream functional assays.

### 2. Predicted Cell-Cell Communication Networks Differ Between Colon Donors

To inform the design of our *in vitro* gut-liver model and derive predictions for immune activation, we explored potential cell–cell interactions using CellPhoneDB^33^. We intergated both immune (Fig.1) and non-immune populations (Fig.S1) identified in tissues. In the colon, we observed strong interaction potential at the trafficking and priming interface, with endothelial cells and fibroblastic reticular cells showing connectivity to multiple neighboring cell types as visualized with chord diagrams (Fig. 2a). Antigen-presenting cells (APCs) such as monocytes, macrophages, dendritic cells, and B cells also demonstrated widespread interaction potential, suggesting active immune surveillance. We found that the number and nature of predicted interactions varied between the two donors, reflecting differences in transcriptional states and unique microenvironments within the colon.

**Figure 2:**
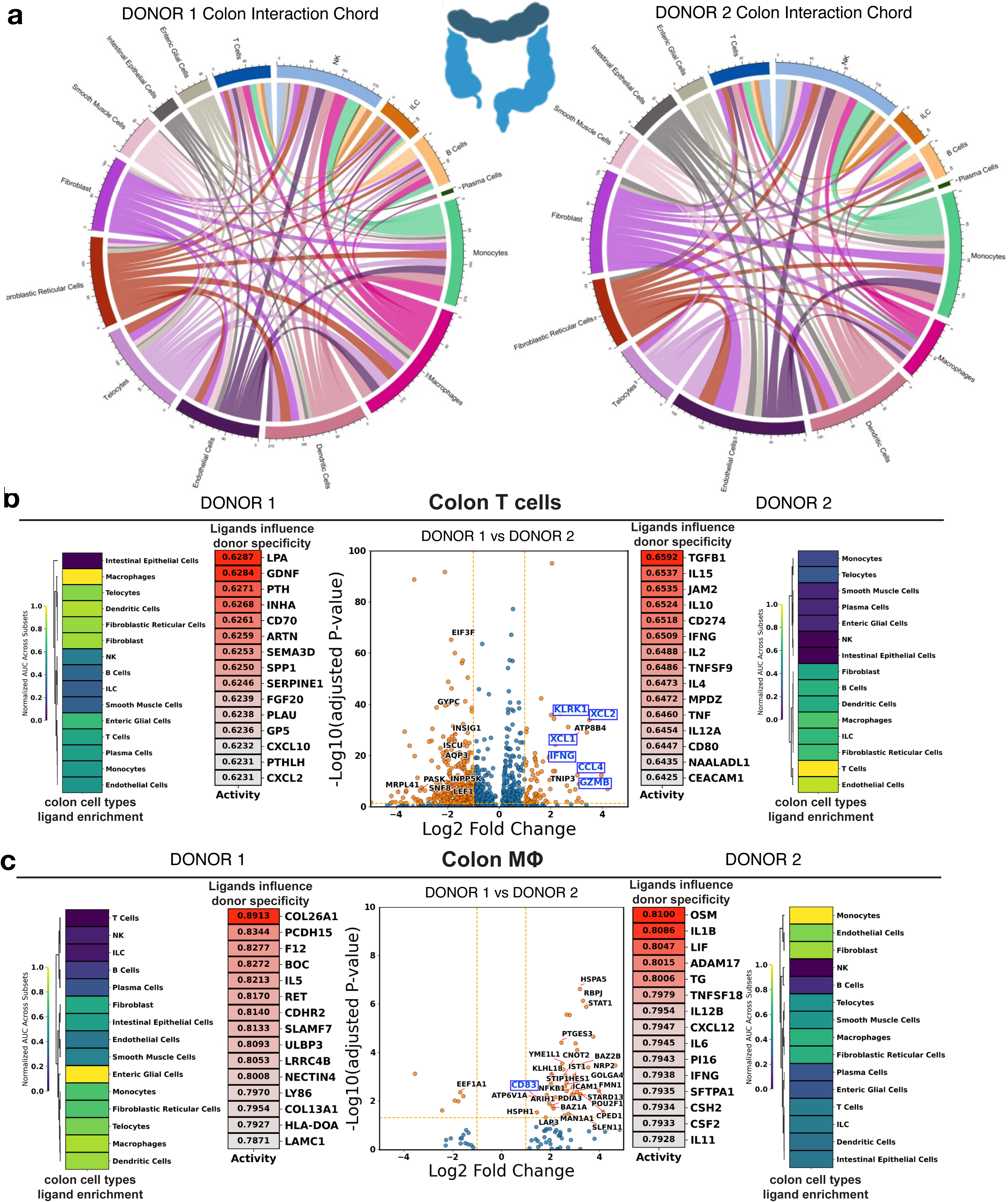
Colon intercellular ligand interactions and differences in donor biology. **a,** graphical representation of the project, approach and its aims. **b,** Uniform Manifold Approximation and Projection (UMAP) embeddings of whole-tissue cells collected in three donor’s colon (left) and liver (right), colored by the donor origin. Only Donor 1 and 2 were used for the model construction. **c,** Top: UMAP embeddings of cells collected in the colon of all donors; Bottom: stacked bar plot shows cell type composition in Donor 1 and Donor 2. **d,** Marker gene expression in all identified colon cell subsets in the sample. The color of each cell in the heatmap denotes the normalized cell type-averaged expression of each marker. NK: natural killer cells; ILC: innate lymphoid cells. **e,** Top: UMAP embeddings of cells collected in the liver of all donors; Bottom: stacked bar plot shows cell type composition in Donor 1 and Donor 2. **f,** Marker gene expression in all identified liver cell subsets in whole-tissue samples. The color of each cell in the heatmap denotes the normalized cell type-averaged expression of each marker. PEC: portal endothelial cells; LSEC: liver sinusoidal endothelial cells.

T cells and macrophages—two key cell types involved in initiating local immune responses— are known for their phenotypic plasticity^35, 36^, shaped by direct cell–cell contact and ligand-mediated signaling. To investigate how these populations differ between donors within the colon, and to understand how local signaling environments contribute to tissue-specific transcriptional profiles, we applied NicheNet^32^, a tool that models ligand–target gene relationships. We supplied NicheNet with differentially upregulated genes from colonic T cells and macrophages in each donor to identify candidate ligands that may drive the observed transcriptional differences. In Donor 1, top ligands influencing T cells—including CD70^37^, CXCL10—were primarily enriched in macrophages and dendritic cells (Fig. 2b, Fig. S2). In contrast, Donor 2’s T cell profile was shaped by ligands such as TGFB1^38^, IL15, IL10 and PD-1 (CD274), which were highly expressed not only in APCs but also in T cells and endothelial cells. For colonic macrophages, BOC and ULBP3 were among the top predicted ligands in Donor 1, with expression predominantly localized to enteric glial cells, highlighting the neural–immune crosstalk previously reported^39^. Meanwhile, in Donor 2, OSM and IL1B were identified as key macrophage-modulating ligands, largely enriched in monocytes and stromal cells (Fig. 2c). Notably, in both donors, macrophages also appeared to influence their own transcriptional programs, suggesting an autocrine feedback loop contributing to niche maintenance.

Based on the described signaling, Donor 1’s colonic T cells exhibit a baseline profile leaning towards activation, driven by APC-derived signals like CD70 and CXCL10. In contrast, Donor 2’s T cells display stronger characteristics of a tolerogenic or regulated state, influenced by inhibitory signals like TGFB1^38^ and PD-1 originating from diverse cell types. This suggests Donor 1 has greater T cell activation readiness, while Donor 2 emphasizes active immune control mechanisms.

### 3. Colon Epithelial Cells Actively Shape the Phenotype and Function of Resident Immune Cells

We established robust colon organoid cultures^29, 40, 41^ from donors 1 and 2 (Fig. 3a) and successfully isolated all CD45^+^ immune populations residing in the intraepithelial space (IEL) and lamina propria (LP)^42^. To assess the impact of the local colonocytes on tissue-resident immune environments, we compared organoid-derived epithelial cells, cultured as a monolayer on membrane inserts, and immune populations to those maintained in co-culture (GUT EPI/IEL/LP) (Fig. 3b). Multiplex cytokine/chemokine assays revealed distinct secretome profiles between conditions, with co-cultures differing from isolated immune or epithelial populations (Fig. 3c,d). PCA visualization, using ClustVis^43^, confirmed distinct clustering based on culture condition (isolated vs. co-culture) and donor origin. Where 51.5% of variability on PC1 explains co-culture versus isolation, while PC2 and 21.3% of variability depend on the donor. Further, a heatmap with clustering of measured cytokines and chemokines across conditions reveals a stark impact of the presence of colonic epithelial cells (EPI) on the cytokine profile of intraepithelial (IEL) and lamina propria (LP) leukocytes (Fig. 3d). Not only do profiles differ among conditions, a number of cytokines can only be reliably detected in co-culture - implying that tissue-resident immune environments act differently in the presence of the colonic epithelium and vice versa. Once combined, Donor 1 co-cultures show a markedly higher concentration of inflammation-associated cytokines (IFNγ, TNFα) as opposed to Donor 2, which shows higher levels of TGFβ-1 and 2. This is in accordance to predictions by NicheNet (Fig. 2b,c).

**Figure 3:**
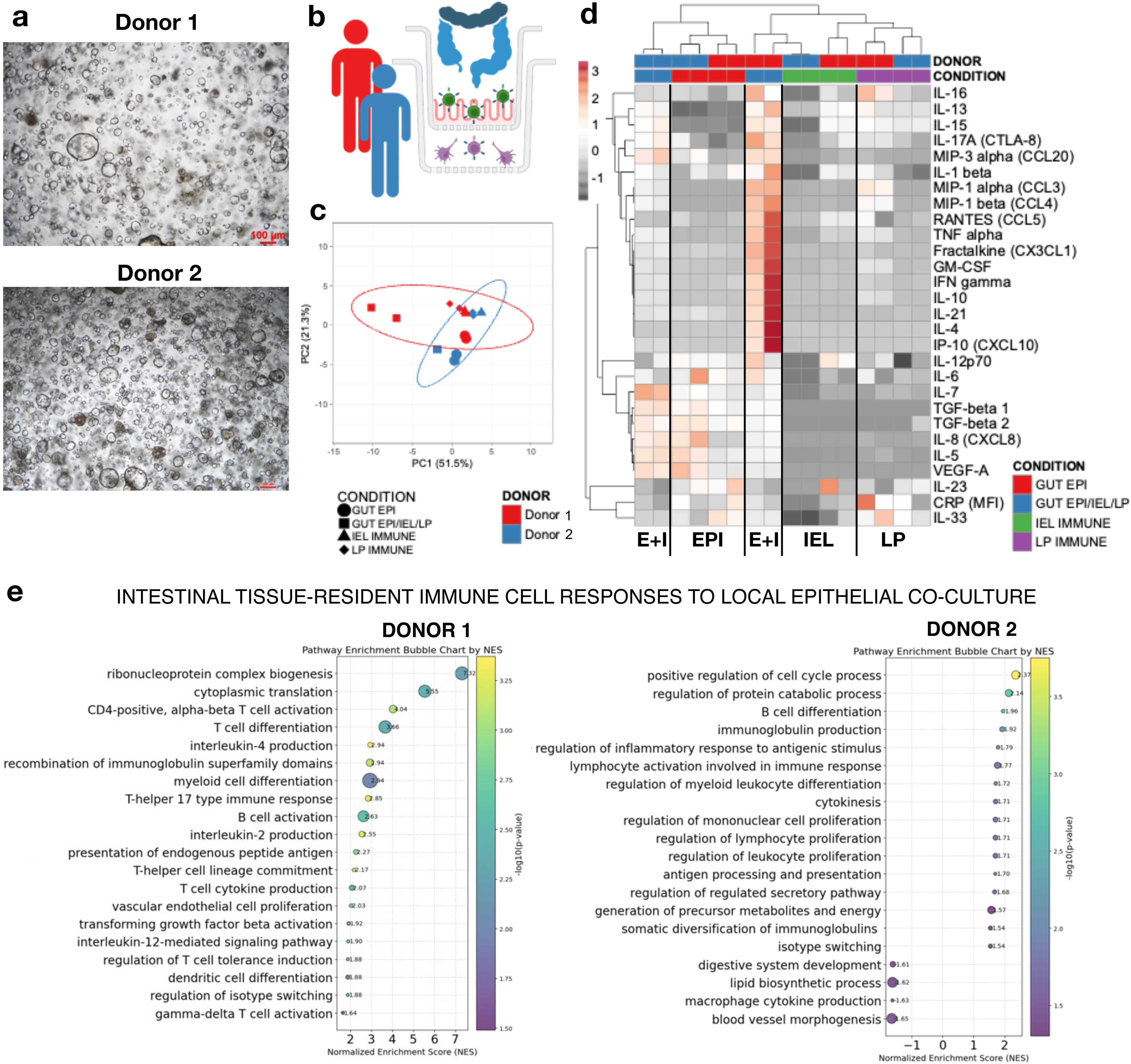
Characterization of an autologous colon co-culture model and its effect on immune cell profiles. **a,** Brightfield images (4x) of established colonic organoids of two donors prior to seeding as monolayers on membrane inserts. **b,** visual representation of the experimental condition where the colonic epithelial cells and intraepithelial immune cells were co-cultured with tissue-resident immune cells derived from the lamina propria or all of them in isolation. **c,** Principal Component Analysis (PCA) performed with ClustVis of measured basolateral cytokines and chemokines released by gut epithelium alone (circular shape), isolated intraepithelial immune cells (triangular shape), lamina propria immune cells (diamond shape) and a co-culture of epithelial cells, intraepithelial leukocytes and lamina propria immune cells (square shape) over 2 days of culture. Variance is explained by PC1 (x=51.5%) and PC2 (y=21.3%). The color red represents values derived from Donor 1 cultures and blue for Donor 2. n=4 per condition and 2 independent experiments. **d,** Heatmap correlation clustering analysis performed with ClustVis for a variety of cytokines and chemokines, measured with Luminex, during a two day cell culture. Rows represent cytokines/chemokines, and columns represent individual samples. Samples (columns) are annotated by condition, where the donors are coded in red (Donor 1) and blue (Donor 2). Further, the gut epithelium cultured alone is in red, intraepithelial leukocytes in green, lamina propria immune cells in purple and a co-culture of epithelial cells, intraepithelial leukocytes and lamina propria immune cells in blue. n=4 per condition and 2 independent experiments. **e,** GSEA pathway enrichment results (normalized enrichment scores (NES), p-values, and gene set sizes) comparing gene expression changes in lamina propria tissue-resident immune cells after co-culture with colon epithelia cells and intraepithelial leukocytes. Left: Donor 1, right: Donor 2. Each graph represents 2 replicates per donor and condition.

We next, performed bulk RNA sequencing on lamina propria tissue resident immune populations, comparing those in isolation to those that were co-cultured with epithelial cells and their matched intraepithelial leukocytes. Gene Set Enrichment Analysis (GSEA; q < 0.05) was used to evaluate donor-specific pathway modulation (Fig.3e). In Donor 1, co-culture significantly enriched pathways related to T cell activation/differentiation (CD4-positive, alpha-beta T cell activation, T-helper 17 type immune response). Conversely, Donor 2 immune cells showed enrichment for B cell differentiation and immunoglobulin production pathways upon co-culture (Fig. 3e). This result is in line with both the NicheNet prediction and secreted cytokine/chemokine profiles. The co-culture results provide functional context to the baseline differences. Donor 1’s immune environment appears primed for strong pro-inflammatory cell-mediated immunity (Th1/Th17) upon epithelial interaction, aligning with baseline activation signals. In contrast, Donor 2’s environment, while showing baseline T cell regulatory signals, is functionally geared towards robust humoral immunity (B cell differentiation and class switching) in the same context, potentially favoring antibody-mediated responses, possibly promoted by baseline TGFβ^44^, over strong T cell inflammation.

### 4. Liver Microenvironment Exhibits Unique Predicted Intercellular Signaling Networks

To explore the donor-specific cellular niches within the liver, we also applied CellPhoneDB and NicheNet to the immune (Fig. 1) and non-immune liver (Fig. S1c,d) scRNA-seq data. While the overall number and pattern of predicted ligand–receptor interactions varied between donors, CellPhoneDB consistently identified macrophages and liver sinusoidal endothelial cells (LSECs) as key hubs of intercellular communication in both individuals (Fig. 4a). This is consistent with their physiological roles—macrophages^45^ as antigen-presenting and cytokine-secreting cells, and LSECs^46^ as gatekeepers of circulatory trafficking. Additional liver-resident cell types, including cholangiocytes, stellate cells, and hepatocytes, were also implicated as important contributors to niche formation. NicheNet analysis further highlighted inter-donor variability in ligand-driven signaling. In Donor 1, T cell-specific transcriptional profiles were most strongly associated with ligands such as CCN4, CD79B, and FGF4, while macrophage-associated ligands included CSF1, CYTL1, and JAM2—all expressed by LSECs (Fig. 4b, c). Conversely, in Donor 2, T cell-influencing ligands like CCL5 and IL10 showed elevated expression in both APCs and cholangiocytes (Fig. 4b). For Donor 2’s liver macrophages, CCL2, OCLN, and CSF1 emerged as top predicted ligands, with expression largely localized to cholangiocytes and APCs (Fig. 4c). Together, these findings suggest that while LSECs are dominant regulators of T cell and macrophage states in Donor 1, cholangiocytes play a more central role in Donor 2. Consistent with our observations in the colon, antigen-presenting cells—including macrophages and dendritic cells—remain key players in shaping immune cell phenotypes within the liver microenvironment.

**Figure 4:**
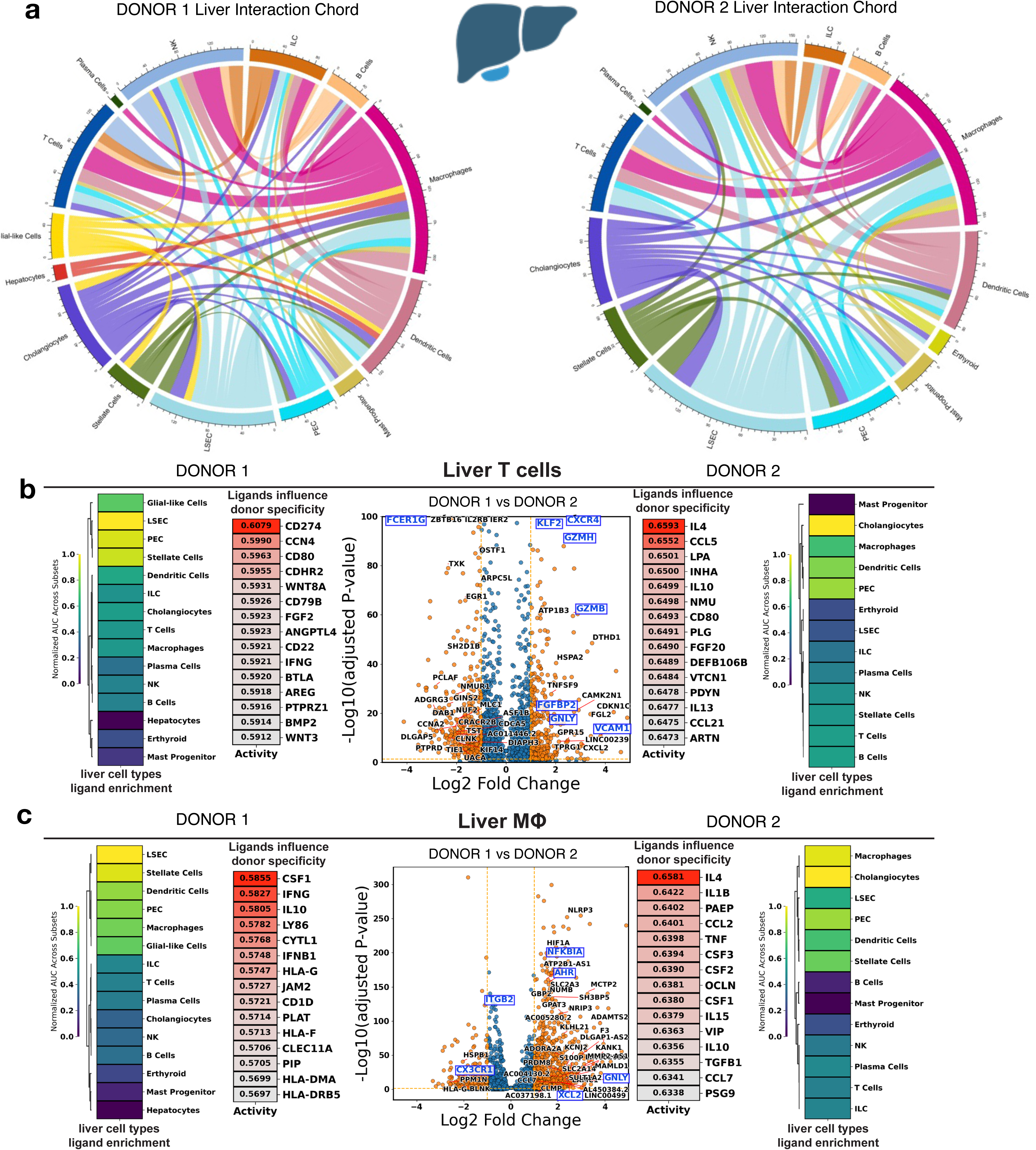
Liver intercellular ligand interactions and differences in donor biology. **a,** Chord diagram showing the number of CellPhoneDB-inferred significant ligand-receptor interactions between each pair of liver cell types in Donor 1(left) and 2(right). **b,c,** Center: Volcano plots showing the differentially expressed genes between **(b)** Donor 1 and 2’s liver T cells **(c)** Donor 1 and 2’s liver Macrophages. X-axis shows the log2 fold change in gene expression between two groups, and the y-axis shows the multi-comparison-corrected P-value of each gene being statistically significant. On each side of the volcano plot is the top 15 (ranked by area under the receiver operating characteristic curve (AUROC)) NicheNet-inferred ligands that shape the differentially upregulated expression (Left to the volcano plot: Donor 1 with respect to Donor 2; Right to the volcano plot: Donor 2 with respect to Donor 1), colored by the auroc value. The enrichment of the NicheNet-inferred donor-specificity-shifting ligands in each donor’s liver cell types were quantified by the area under the recovery curve (AUC) against the top 30 (ranked by area under the receiver operating characteristic curve (AUROC), AUROC > 0.5) NicheNet-inferred ligands. Results are visualized by heatmap, where the color represents the enrichment min-max normalized across cell types.

Contrasting with their respective gut profiles, the baseline liver immune analysis reveals distinct donor environments. Donor 1’s liver niche appears more homeostatic and focused on tissue interaction/remodeling via LSEC signaling (CSF1, CCN4, FGF4), differing markedly from the activation-prone T cell signature seen in their gut. Conversely, Donor 2’s liver shows significant signaling involving cholangiocytes and APCs, resulting in a dynamically balanced baseline; notably, this liver environment features concurrent signals for potential immune recruitment/inflammation (CCL5, CCL2) alongside strong regulation (IL10), suggesting a different immune posture and potentially greater baseline management of inflammatory signals compared to the predominantly T cell-regulatory profile and humoral focus observed in Donor 2’s gut.

### 5. Hepatocytes Influence the Transcriptional State and Secretome of Resident Liver Immune Cells

To create the local liver MPS model we cultured isolated primary hepatocytes in a sandwich configuration that supported hepatocyte viability and polarization as determined by expression of Zona Occludens-1 (ZO-1) proteins on points of contact (Fig. 5a)^47–49^. We assessed the impact of polarized hepatocytes grown on inserts on tissue-resident immune environments bellow as opposed to each niche cultured in isolation (Fig. 5b). PCA visualization showed distinct clustering based on culture condition (isolated (LIV IMMUNE, LIVER HEP) vs. co-culture (LIVER HEP/IMMUNE)) and donor origin. 65.5% of variability on PC1 explains co-culture versus isolation, while PC2 and 14.1% of variability depend on the donor (Fig. 5c). Heatmap with clustering of measured cytokines and chemokines across conditions confirmed a distinct impact of the presence of hepatocytes (LIVER HEP) on the cytokine profile of liver-resident immune populations (LIV IMMUNE) (Fig. 5d). As in the case of colonic-immune cultures, a number of cytokines was only measured under co-culture condition emphasizing the importance of tissue-resident environments as opposed to studies with hepatocytes only. In isolation, cultured hepatocytes and hepatic immune cells, are virtually indistinguishable by donor, however, when combined Donor 2 co-cultures lead to a significantly different cytokine/chemokine environment as Donor 1. In concurrence to NicheNet predictions, Donor 2 co-cultures showed higher concentrations of cytokines associated with inflammation (TNFα, IFNγ, IL-17) and immune recruitment (CCL5, IP-10).

**Figure 5:**
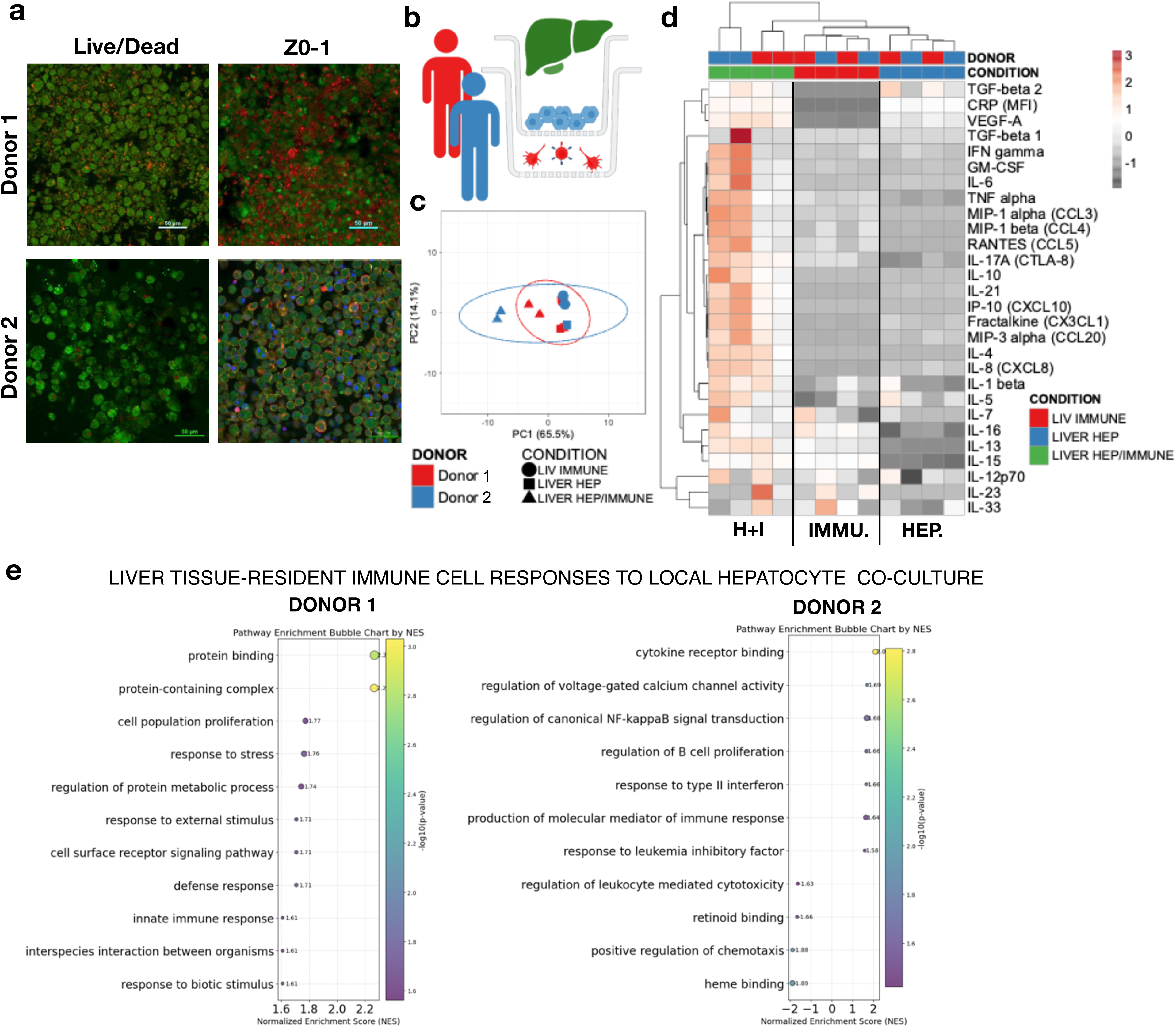
Characterization of an autologous liver co-culture model and its effect on immune cell profiles. **a,** Confocal images (10x) of established primary hepatocyte sandwich cultures on membrane inserts of Donor 1 (top) and 2 (bottom) after 4 days in culture. Left column shows viable cells (green) and non-viable cells (red). Right column shows cells stained with DAPI (nucleus – blue), Phalloidin (F-Actin – green) and Z0-1 (canaliculi – red). **b,** visual representation of the experimental condition where primary hepatocytes were cultured in a sandwich configuration in co-culture with tissue-resident immune cells or both in isolation. **c,** Principal Component Analysis (PCA) performed with ClustVis of measured basolateral cytokines and chemokines released by hepatocytes alone (square shape), isolated liver immune cells (circular shape), and a co-culture of hepatocytes and tissue-resident liver immune cells (triangular shape) over 2 days of culture. Variance is explained by PC1 (x=65.5%) and PC2 (y=14.1%). The color red represents values derived from Donor 1 cultures and blue for Donor 2. n=4 per condition and 2 independent experiments. **d,** Heatmap correlation clustering analysis performed with ClustVis for a variety of cytokines and chemokines, measured with Luminex, during a two day cell culture. Rows represent cytokines/chemokines, and columns represent individual samples. Samples (columns) are annotated by condition, where the donors are coded in red (Donor 1) and blue (Donor 2). Further, the hepatocytes cultured alone are in blue, tissue-resident immune cells in red, and a co-culture of hepatocytes and liver tissue-resident immune cells in green. n = 4 per condition and 2 independent experiments. **e,** GSEA Pathway enrichment results (normalized enrichment scores (NES), p-values, and gene set sizes) comparing gene expression changes in tissue-resident liver immune cells after co-culture with hepatocytes. Left: Donor 1, right: Donor 2. Each graph sents 2 replicates per donor and condition.

Next, we investigated how the co-culture of liver-resident immune populations with hepatocytes influences their gene expression (Fig.5e). GSEA (q < 0.05) revealed pathways related to stress response, protein metabolism, and innate immunity in Donor 1 immune cells. In Donor 2, co-culture enriched NF-kappaB signaling, B cell proliferation, and type II interferon response pathways in agreement with cytokine measurements (Fig. 5e).

### 6. Fluidic Integration of the Colon and Liver MPS Leads to Significantly Altered Behavior and is Consistent Across Donors

After the characterization of colonic and liver immune co-cultures individually, we asked how fluidic interaction between these two organ systems affects their biology and if the effects of the interaction differ between donors (Fig. 6a,b,c). Colon monolayers with intraepithelial leukocytes were grown in 24-well inserts. These were then placed into larger 12-well inserts containing lamina propria immune cells. Similarly, hepatocytes were grown in a sandwich configuration in 24-well inserts and placed into 12-well inserts with liver-resident immune cells. Gut and liver immune co-cultures were positioned into a two-compartment Multiorgan Tissue Interaction Vessel (MOTIVE-2) (Fig.6b). The MOTIVE-2 vessels were connected to peristaltic pumps on the battery-driven MOTIVE platform. Each platform fits three vessels and allows for various perfusion rates. The controller is operated either manually or via Bluetooth. MOTIVE-2 vessels were 3D printed in-house and are made of biocompatible Digital ABS Plus material.

**Figure 6:**
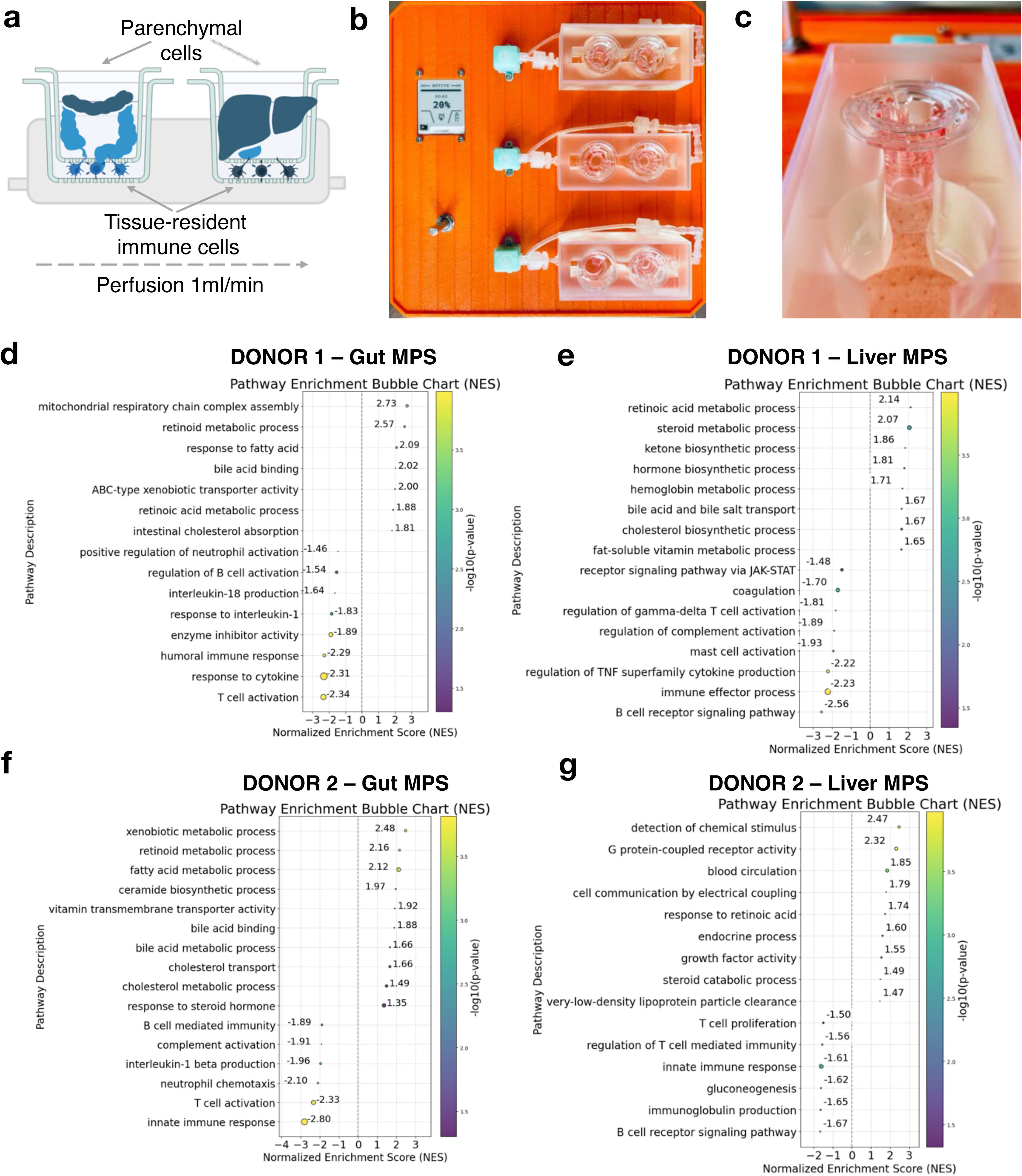
Establishing an autologous Gut-Liver interaction MPS with respective tissue-resident immune environments. **a,** Visual representation of the MPS system and experimental set-up. The colon co-culture system was created by culturing colon epithelial cells and intraepithelial leukocytes in a 24-well membrane insert and placing it on top of a 12-well insert containing lamina propria tissue-resident leukocytes. The liver co-cultures were established by seeding primary human hepatocytes in a sandwich configuration in 24-well inserts and placed into a larger 12-well insert, which contained liver tissue-resident immune cells. Both compartments were fluidically connected in a two-well MultiOrgan Tissue Interaction Vessel (MOTIVE-2) at a perfusion rate of 1ml/min. Fluid dynamics and operation in the MOTIVE-2 vessels were controlled by a custom-built MOTIVE platform. **b,** MOTIVE platform with a controller and display controlling the operation of three peristaltic pumps that perfuse MOTIVE-2 vessels with universal media. **c,** Cross-sectional view of the MOTIVE-2 vessel and one of the compartments containing a 24-well/12-well sandwich set up. **d-g,** GSEA Pathway enrichment results (normalized enrichment scores (NES), p-values, and gene set sizes) comparing gene expression changes between colon epithelial/immune co-cultures in isolation versus interaction with co-cultures of hepatocytes and their respective immune cells **(d – Donor 1, f-Donor 2)**. Vice-versa GSEA Pathway enrichments were obtained from hepatocytes cultured with their respective immune cells in isolation or interaction with intestinal/immune co-cultures **(e – Donor 1, g-Donor 2)**. Each graph represents 2 replicates per donor and condition.

One of the first control experiments was to compare multiplexed cytokine data in common medium during gut-liver interaction in either the full immunocompetent set-up or interacting epithelium and hepatocytes in the complete absence of any immune cells. As expected the addition of their respective immune-resident environments led to a strong increase in cytokine/chemokine concentrations. Donor 1 common media was defined by higher IFNγ, IL23 and IL1 while Donor 2 had more IL6, IL-17A and TNFα (Fig. S3). These differences were relatively minor and only 18% variance (PC2) was explained depending on the donor. We harvested the interacting tissues after two days of perfusion with common media and determined gene expression changes (Fig. 6d-g). Further analysis isolating the interaction effect (interaction vs. isolation co-culture) showed that interaction had a surprisingly similar effect on tissues of both donors. In particular, pathways enriched in intestinal tissues during interaction confirm expression of genes associated with a response to liver-derived factors such as retinoid^50^ and bile acid binding^51, 52^. Moreover, the metabolism of fatty acids and cholesterol transport are noted in both donors. Simultaneously, interaction led to the reduced expression of pathways associated with inflammation and immune activation (T cell activation, B cell immunity, IL1 signaling).

In strong concurrence with these findings, interacting liver/immune tissue showed increased expression of pathways associated with retinoic acid metabolism, steroid metabolism, cholesterol synthesis and bile acid transport. (Fig. 6e, g) while simultaneously downregulating pathways associated with immune cell activation such as T cell mediated immunity, immunoglobulin production and innate immune responses. Interestingly, a previous MPS model of the gut-liver axis that was established through co-culture of gut epithelial cells with PBMC-derived macrophages and DC as well as primary hepatocytes and Kupffer cells, also showed the interaction of the two organ systems to increase liver metabolism and to reduce immune activation^29^.

The data suggests that core physiological crosstalk mechanisms might be conserved despite baseline immune variations, or possibly that the in vitro environment drives this convergence.

### 7. Functional Responses of Colon Tissue Model to Challenges are Donor- and Stimulus-Specific

Finally, we were interested in understanding how the reconstructed autologous gut-liver-immune models respond to intestinal immune perturbation in the individual donors during the very first days of immune activation to capture early signaling cascades. We challenged the integrated colon model with either the 1.) lipopolysaccharides (LPS) that are bacterial membrane components and recognized by Toll-like Receptor 4^53^; 2.) Polyinosinic:polycytidylic acid (Poly I:C), a synthetic analog of double-stranded viral RNA and recognized via Toll-like Receptor 3^54^ or 3) 5-(2-oxopropylideneamino)-6-d-ribitylaminouracil (5-OP-RU), a known antigen recognized by Mucosa Associated Invariant T (MAIT) cells who represent a high percentage of liver T cells^5, 26^. 5-OP-RU is a derivative of intermediates produced during bacterial riboflavin biosynthesis (Fig.7a).

After activation and two days of interaction in the MOTIVE-2 vessels, multiplexed cytokines and chemokines were analyzed in the apical gut compartment (Fig.7b). Most notably, Poly I:C, as the most potent of the three immune activators, led in both donors to a strong response with significantly induced production of the interferon-induced IP-10, and the cytokines IL4, IL6 and IL-21. On the other hand both LPS and 5-OP-RU were in their action less universal and dampened compared to Poly I:C. LPS treatment led to significant production of CX3CL1, TNFα and modestly increased IL-6 in Donor 1 but not Donor 2. No significant cytokine/chemokine changes were observed in the apical colon compartment after 5-OP-RU treatment. Both TLR 3 and 4 are abundantly expressed among immune cells and epithelial cells in the colon/IEL section of the gut MPS^55^, while 5-OP-RU is a MAIT cell-specific antigen and thus less likely to lead to expression of inflammatory mediators directly in the lumen. Nonetheless, all three immune activators led to a marked reduction of gut barrier function as determined with measurements of the trans epithelial electrical resistance (TEER) in both donors (Fig. 7c). Despite the reduction in TEER the barriers remained intact (Fig. S4) at the end of the experiment, where the lowest TEER measurement across all conditions was still higher than 800 Ohms.

**Figure 7:**
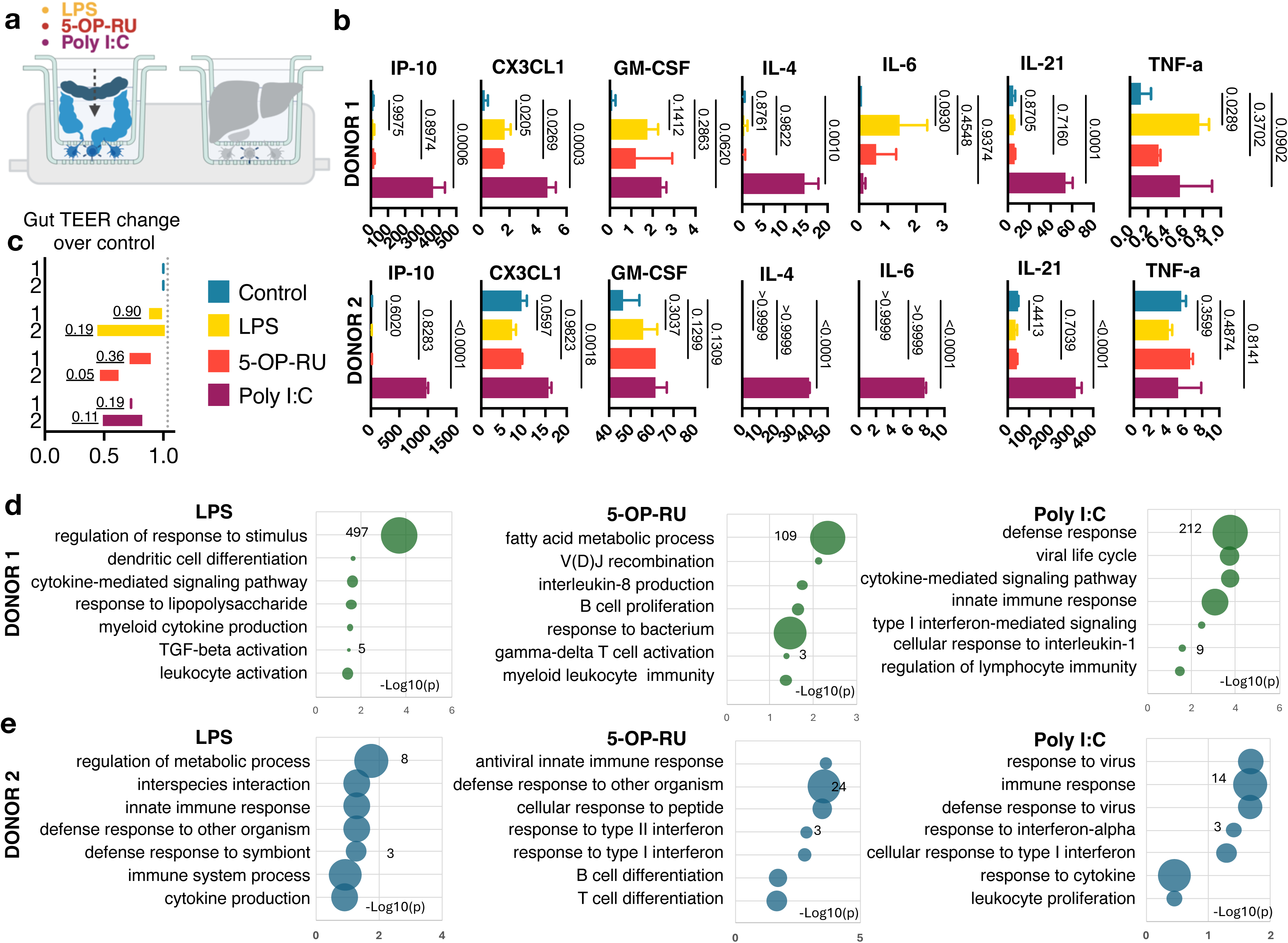
Colonic immune responses to a challenge with LPS, 5-OP-RU and Poly I:C during gut-liver-immune interaction. **a,** Visual representation of experimental design and measured parameters, where a colonic co-culture of epithelial cells and their tissue-resident immune cells was fluidically connected to a liver model composed of hepatocytes and liver resident immune populations over two days and an intestinal challenge with LPS, 5-OP-RU, or Poly I:C. **b,** Cytokine and chemokine concentrations measured via Luminex in the apical media of colonic co-cultures during interaction with a hepatic/immune model and during an immune activation challenge with either LPS (yellow), 5-OP-RU (orange) or Poly I:C (purple) and control without activation (blue). Each graph represents 2 replicates per donor and condition. Post-hoc pairwise comparisons against the control were conducted following ANOVA using Fisher’s Least Significant Difference (LSD) test to determine statistical significance (p values annotated on bar graphs). **c,** Changes in Trans-epithelial electrical resistance (TEER) of Donor 1 and Donor 2 colonic monolayers in co-cultures while challenged with either LPS (yellow), 5-OP-RU (orange) or Poly I:C (purple). Impacts on TEER are expressed as fold changes over the control. Post-hoc pairwise comparisons against the control were conducted using Fisher’s Least Significant Difference (LSD) test to determine statistical significance (p values annotated on bar graphs).**d-e**, GSEA Pathway enrichment results (normalized enrichment scores (NES), p-values, and gene set sizes) comparing gene expression changes in colonic epithelial/immune co-cultures during interaction with the liver compartment and during an immune activation challenge with either LPS, 5-OP-RU or Poly I:C. Donor 1 responses are presented under **(d)** and Donor 2 under **(e).** Each graph represents 2 replicates per donor and condition.

In agreement with the reduction in TEER, GSEA analysis of differentially expressed genes across conditions and donors confirmed the nature of individual immune activators and their elicited reponse in the gut/immune MPS (Fig.7d,e). Predictably, pathways associated with a response to LPS and innate immune activation, in particular in Donor 1, were identified in the LPS-treated group in both donors. Both donors also showed a robust enrichment in pathways associated with viral control and interferon-mediated immunity in the Poly I:C group. Interestingly, 5-OP-RU appears to have led to B cell activation and VDJ recombination as well as T cell differentiation in both donors. Neither LPS or Poly I:C seemed to influence adaptive immune responses at this early stage of exposure.

### 8. Liver Tissue Model Displays Donor-Dependent Functional Responses to Challenges

At the same time, we evaluated immunological responses in the liver MPS compartment following the intestinal challaneg with LPS, 5-OP-RU and Poly I:C (Fig.8a). Given the reduced baseline state of immune activation in the liver MPS during interaction (Fig.6) and short interaction period to catch the earliest responses, only modest changes were observed.

Nonetheless, the findings indicate donor-specific differences in the early response to intestinal immune activation. Multiplexed cytokine/chemokine data shows higher mean values of the IP-10, CX3CL1, IFNγ and IL-21 in Donor 1 post intestinal LPS challenge albeit not significant. On the other hand, higher mean values of IP-10, CX3CL1, IFNγ, IL-8, IL-21 and the fibrotic liver marker FGF-21^56^ were measured in Donor 2 during the intestinal challenge with 5-OP-RU and Poly I:C (Fig.8b). Albumin production remained relatively stable in both donors following colonic challenges with all three activators as compared to the controls (Fig.8c). Frequently, gene expression changes precede measurable changes in cytokine/chemokine production (Fig.8d,e). In the case of Donor 1, GSEA analysis (q < 0.05) revealed no significant differences in the 5-OP-RU treated group. However, early adaptation to LPS treatment was evident through increased transmembrane transport and to Poly I:C via increased interferon, autophagy and angiogenesis signaling. Similar transcriptomic responses were noted in Donor 2 following the Poly I:C challenge. But in contrast to Donor 1, no significant changes were observed in the LPS-treated group, while intestinal exposure to 5-OP-RU led to an increased response to external stimulus, metabolic oxoacid adaptations, and increased angiogenesis. A possible explanation of a greater response to 5-OP-RU by Donor 2 could be the increased proportion of CD8 TCRαβ MAIT cells^57^ (Fig. S5) found among hepatic T cells as compared to Donor 1 (Fig. 8f,g). Similarly, Donor 1 single-cell RNA seq data has shown a higher percentage of macrophages in the tissue-resident immune cell pool, which could be one of the possible reasons for greater susceptibility of Donor 1 to an intestinal challenge with LPS^58^ (Fig. 8g)

**Figure 8:**
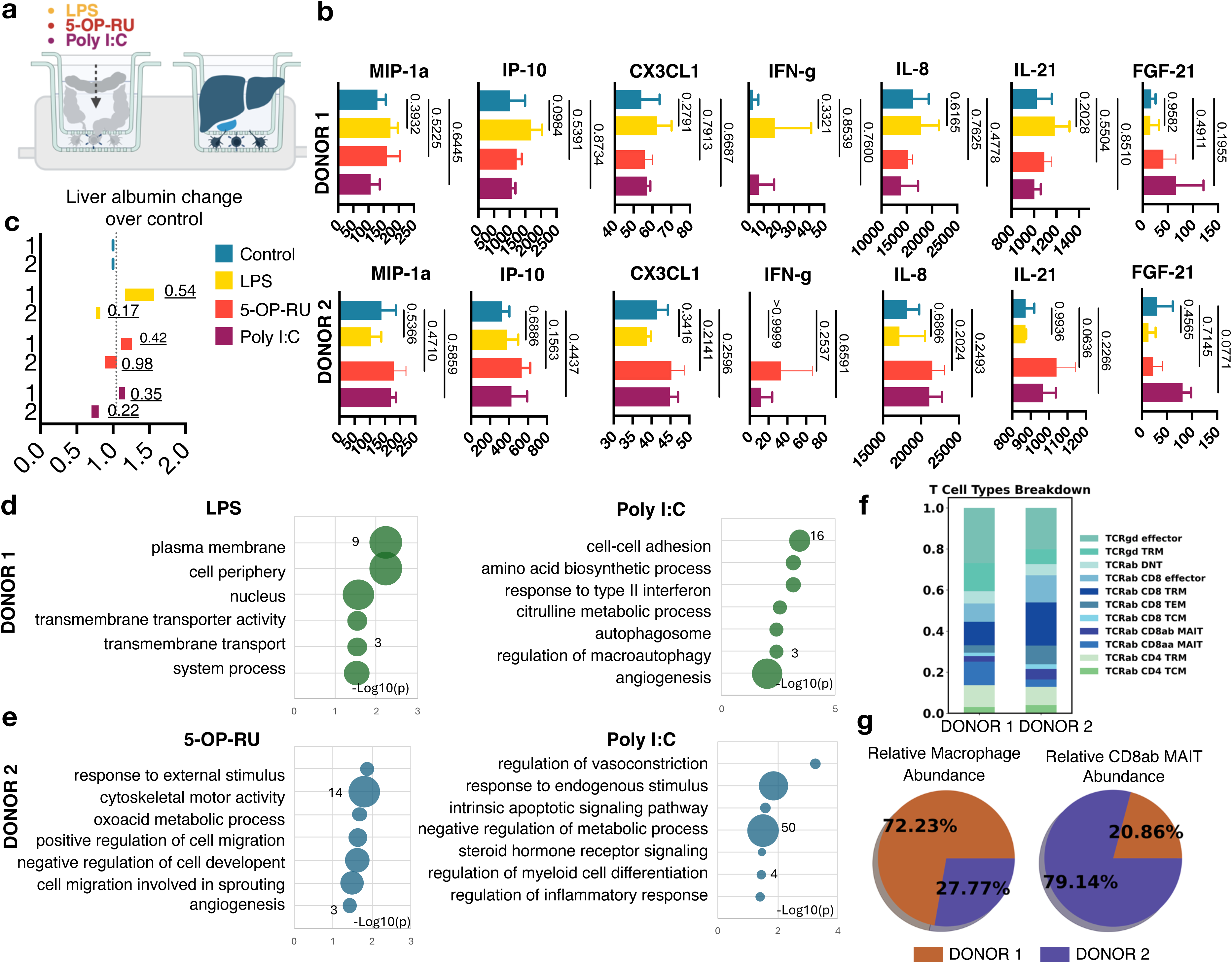
Hepatic immune responses to an intestinal challenge with LPS, 5-OP-RU, and Poly I:C during gut-liver-immune interaction. **a,** Visual representation of experimental design and measured parameters, where a colonic co-culture of epithelial cells and their tissue-resident immune cells was fluidically connected to a liver model composed of hepatocytes and liver resident immune populations over two days and an intestinal challenge with LPS, 5-OP-RU, or Poly I:C. **b,** Cytokine and chemokine concentrations measured via Luminex in the apical media of hepatocyte/immune co-cultures during interaction with a colonic/immune model and during an immune activation challenge of the gut MPS with either LPS (yellow), 5-OP-RU (orange) or Poly I:C (purple) and control without activation (blue). Each graph represents 2 replicates per donor and condition. Post-hoc pairwise comparisons against the control were conducted following ANOVA using Fisher’s Least significant Difference (LSD) test to determine statistical significance (p values annotated on bar graphs). **c,** Changes in albumin secretion of Donor 1 and Donor 2 hepatic co-cultures during intestinal challenge with either LPS (yellow), 5-OP-RU (orange) or Poly I:C (purple). Impacts on albumin secretion are expressed as fold changes over the control. Post-hoc pairwise comparisons against the control were conducted using Fisher’s Least Significant Difference (LSD) test to determine statistical significance (p values annotated on bar graphs).**d-e**, GSEA Pathway enrichment results (normalized enrichment scores (NES), p-values, and gene set sizes) comparing gene expression changes in hepatic/immune co-cultures during interaction with the gut compartment and during an intestinal immune activation challenge with either LPS, 5-OP-RU or Poly I:C. Donor 1 responses are presented under **(d)** and Donor 2 under **(e).** Each graph represents 2 replicates per donor and condition. **f,** Stacked bar plot showing the T cell subsets composition in Donor 1 and 2’s primary liver tissue. **g,** Pie chart showing the abundance of macrophage (left) and CD8αβ MAIT cells (right) in donors’ primary liver tissue.

Cumulatively both differences in ligand interactions and cellular composition of the immune niche may contribute to donor-specific responses in the immune challenge of the gut-liver-immune axis.

## Discussion

The present study establishes an autologous, immunocompetent gut–liver MPS that is anchored in single-cell provenance and validated by multiomic read-outs. By integrating donor-resolved atlases of the human colon and liver with bespoke parenchyma-immune co-cultures and a perfused MOTIVE platform, we uncover how individual tissue-resident immune landscapes translate into distinct, yet partially convergent, functional states. To our knowledge, this is the first demonstration that faithfully preserves donor-matched lamina-propria, intra-epithelial and hepatic immune niches within a dual-organ circuit, enabling interrogation of inter-organ immune crosstalk.

Our single-cell analysis confirmed the establishment of diverse cellular ecosystems within the MPS, consistent with recent human tissue atlases^2–4^, but crucially highlighted the significant inter-donor variability inherent in healthy individuals^18^. Computational modeling predicted distinct immune ‘set points’ for each donor and organ, driven by different signaling hubs and ligand environments (e.g., pro-activation vs. regulatory signals^32, 33, 37–39^). This baseline characterization emphasizes the necessity of incorporating primary cells and donor variability to truly model human immunological diversity.

Partial validation through co-culture experiments underscored the importance of parenchymal-immune interactions, often overlooked in simpler models. The presence of epithelial cells or hepatocytes significantly altered immune cell secretomes and transcriptional states, largely aligning with the computationally predicted functional polarization (e.g., Th1/17 vs. B-cell responses in the colon^44^). These findings validate both the predictive power of tools like NicheNet and the critical role of the tissue microenvironment in shaping resident immune function, supported by established culture methods^40, 41, 47^.

Strikingly, despite baseline donor differences, fluidic connection of the gut and liver modules induced conserved responses across individuals. Both donors exhibited metabolic adjustments reflecting known gut-liver biochemical communication^50–52^ and, counterintuitively, a significant dampening of immune activation pathways. This convergence towards a homeostatic, potentially tolerogenic state upon interaction, consistent with prior MPS work^29^, suggests dominant physiological crosstalk mechanisms may regulate systemic immunity, overriding some baseline individual variations in the healthy state. Capturing such emergent systemic properties is a key advantage of multi-organ MPS.

The platform’s utility was further demonstrated by its response to intestinal challenges. The system differentiated between various immune stimuli^5, 26, 53, 54^ and, importantly, displayed donor-specific functional outcomes that could be rationalized by the initial baseline characterization – linking specific cell proportions (e.g., liver macrophages or MAIT cells^57, 58^) to differential susceptibility or response pathways. This capacity to connect inherent donor features to functional responses represents a critical step towards predictive modeling of individual human variation.

Central to the model is the MOTIVE-2 platform, whose features—including user accessibility, battery-driven operation enabling transport under perfusion—facilitate the execution and observation of complex, interacting co-culture experiments. The platform captures the interaction between parenchymal cells and complex tissue-resident immune populations across the human gut-liver axis, revealing insights into baseline heterogeneity, homeostatic crosstalk, and innate/MAIT responses. This focus on multi-lineage TRI immunity and inter-organ communication complements other recent developments^31^. While such systems provide deep insights into specific lymphocyte subsets and adaptive immunity within a single tissue, our multi-organ model offers a distinct advantage for studying the broader TRI landscape, innate-adaptive interplay, systemic metabolic-immune crosstalk, and how baseline variability across organs influences initial responses to challenges originating in the gut. By overcoming limitations of animal models and simpler in vitro cultures, this integrated approach provides a new tool for dissecting human immunology, modeling disease, and improving preclinical drug evaluation.

Nevertheless, limitations exist, including the small donor number, the unavoidable simplification compared to in vivo complexity (e.g., lacking microbiome, full stromal components, lymph node), and the short experimental duration. A limitation of the present study is the use of a small donor number. While this work provides an unprecedentedly deep, multi-omic proof-of-concept of how individual immune set-points translate to function, future studies with a larger cohort of donors will be necessary to establish broader generalizations across the population. However, the ability to rationalize the distinct functional outcomes in these two donors based on their initial cellular profiles provides a strong framework for interpreting human immune diversity in future, scaled-up studies. Further, the ligand-receptor networks are in silico predictions awaiting perturbational validation. Longer perfusion windows and untargeted metabolomics will be required to link transcriptomics to metabolite flux.

In conclusion, this immunocompetent, donor-specific, human gut-liver MPS provides unique insights into tissue homeostasis, immune heterogeneity, and inter-organ crosstalk. It represents a platform for advancing our understanding of human physiology and offers significant potential for more predictive disease modeling and preclinical evaluation of therapeutics. Lastly, it represents a stepping stone in expanding donor-matched models to additional organ systems and the integration of secondary and tertiary lymphoid structures.

## Methods

### HUMAN SUBJECTS

Human colon tissue, liver tissue, and blood samples were purchased from LifeNet Health LifeSciences. The first non-diseased donor (Donor 1) was a Caucasian female aged 28 with BMI:20. The second non-diseased donor (Donor 2) was a Caucasian male aged 43 with BMI:24.3. The tissue was transported in the University of Wisconsin perfusion media on ice. It arrived in our facilities in under 24 hours of cold ischemia time and under 6 hours of warm ischemia time.

### PROCESSING OF TISSUES AND PREPARATION OF CO-CULTURES

#### Isolation of human colonic crypts from fresh colon tissue

Tissue was processed based on a previously established protocol^42^. In brief, upon arrival, colon tissue was divided into three equal pieces using a scalpel and sharp scissors at room temperature. Colon tissue was washed with pre-rinse buffer containing Hank’s 1X Balanced Salt Solutions (HBSS) without calcium and magnesium (Cytiva, Cat no: SH30588.01) and penicillin-streptomycin-amphotericin B (PSA) (Lonza, Cat no: 17-745E). First, adipose tissue was removed, and then the epithelium was separated from the muscularis externa with forceps. Each trimmed colon tissue was cut into small pieces with scissors and placed into small tissue culture dishes (21.2 cm^2^) (Olympus Cat No: 25-260). Tissues were minced into fine pieces with an approximate size of 5mm using the McIlwain Tissue Chopper (Cavey Laboratory Engineering Co. Ltd). Meanwhile, freezing media containing 10% DMSO (Tocris, Cat no: 3176) and 90% HI-FBS (Cytiva HyClone, Cat no: SH30910.03) was prepared. Each chopped tissue piece was mixed with freezing media and distributed into cryopreservation tubes placed in a freezing container, frozen at −80°C overnight, and transferred to −196°C for further use. On the day of isolation of human colonic crypts from cryopreserved colon tissue, we prepared i) the rinse/thawing buffer containing RPMI-1640 (Gibco, Cat no: 11875-093), 5% HI-FBS (Cytiva HyClone, Cat no: SH30910.03) and 1% Penicillin-Streptomycin (PS) (Gibco, Cat no: 15-140-122), ii) pre-digestion buffer with 5mM EDTA ((Invitrogen; Cat no: 15575-038), 1 mM DTT (Teknova; Cat no: D9750), 10 mM HEPES (Gibco, Cat no: 15630-080), 1% PSA (Lonza, Cat no: 17-745E), 0.1 % BSA (Fisher, Cat no: BP1600-100) in 1x HBSS without calcium and magnesium (Cytiva, Cat no: SH30588.01). All buffers were warmed at 37°C.

One vial of cryopreserved colon tissue pieces was thawed swiftly at 37°C, washed with the rinse/thawing buffer, and transferred to a 50 ml tube containing the rinse/thawing buffer. Then, it was centrifuged at 400xG for 10 minutes. The supernatant was discarded, and the pellet was resuspended with 15 ml of prewarmed predigesting buffer by inversion. Then, it was incubated horizontally for 20 min at 37°C under continuous rotation (140 RPM) using an incubating shaker (Thermo Scientific, MaxQ 420HP). After that, the tissue pieces waited to settle to the bottom of the 50-ml tube, and then the supernatant containing the stripped epithelium was transferred into a new 50-ml conical tube placed on ice using a pre-wetted 25ml serological pipette. The colonic crypts were observed under microscopy. Another 15ml of fresh prewarmed predigesting buffer was added to the colon tissue pieces and incubated horizontally for 20 min at 37°C under continuous rotation (140 RPM) using an incubating shaker (Thermo Scientific, MaxQ 420HP). After the second round of epithelial stripping, the tube was mixed gently for 10 seconds using a vortex and then waited to be settled at the bottom of the 50-ml conical tube. The second supernatant containing the stripped epithelium was collected and combined with the first round into a 50ml conical tube placed on ice using a pre-wetted 25 ml serological pipette. The colonic crypts were checked again under the microscope. The 100μm cell strainer was washed with 5-10ml of rinse buffer, and the combined supernatant was passed through a 100μm cell strainer placed on a new 50-ml conical tube. 15 ml of ice-cold Rinse buffer was added to the tissue pellet, then centrifuged at 400xG for 10 minutes. The supernatant was transferred to a tube with a 100μm cell strainer and combined with previous supernatants. This combined cell suspension containing stripped epithelium was centrifuged at 500g for 10 minutes. The pellet was resuspended with rinse buffer. Total cell number and cell viability were determined with an automated cell counter (Denovix, CellDrop FL) using trypan blue staining.

#### Colon organoid culture

Organoid cultures were initiated as previously described^29^. Before starting the procedure, the basement membrane Matrigel® Matrix (Corning, Cat no: 356231) was thawed overnight at 4°C. 24 well flat-bottom tissue culture plates (GenClone, Cat no: 25-107) were prewarmed at 37°C for at least 2 hours before use for organoid culture. 250.000 cells per dome were slowly and thoroughly mixed with 50μl of Matrigel® Matrix (Corning, Cat no: 356231) per dome on ice using a pre-wetted pipette tip. 50μL droplets containing Matrigel®-crypt suspension were seeded as domes onto the wells of a 24-well tissue culture-treated plate using a pre-wetted pipette tip. The plate was carefully transferred to a 37°C incubator and incubated at 37°C for 15-20 minutes to allow the domes to solidify. In the meantime, organoid culture media was prepared as follows: L-WRN-conditioned media supplemented with 2mM Glutamax (Gibco, Cat no: 35050-061), 10mM HEPES (Gibco; Cat no: 15630-080), 100 units/ml Pen/Strep (Gibco, Cat no: 15-140-122), B-27 supplement (Gibco, Cat no: 17504-001), 10 mM nicotinamide (Sigma-Aldrich, Cat. no. N0636-100g), N-2 supplement (Gibco, Cat no: 17502-048), 500 μM N-acetyl cysteine (Sigma-Aldrich, Cat no: A7250-5g), 500 nM A83-01 (Tocris, Cat no: 2939), 10μM Y-27632 Dihydrochloride (Biogems, Cat no: 1293823-10mg), 10 μM SB 202190 (Biogems, Cat no: 1523072-25mg), 50 ng ml^−1^ recombinant human EGF (Gibco, Cat no: PHG0313), 10 nM human [Leu15]-gastrin I (Anaspec, Cat no: AS-64149), 5 nM prostaglandin E2 (Biogems, Cat no: 3632464-10mg). 600μl of organoid culture media was added to each well and incubated at 37°C and 5% CO_2_. Media was refreshed every 3-4 days. After 7-14 days, organoids were expanded by incubating in Cell Recovery Solution (Corning, Cat no: 354253) for 45-60 min at 4°C, followed by mechanical dissociation and replated in fresh Matrigel® (Corning, Cat no: 356231) at a 1:3 ratio. Colon organoids were passaged every 7-10 days in fresh Matrigel® (Corning, Cat no: 356231) at a 1:3 ratio using Cell Recovery Solution (Corning, cat. no. 354253) for 45-60 min at 4°C and mechanical dissociation.

#### Seeding colon epithelial monolayers on membrane inserts

Before starting the procedure, 24-well 0.4 μm PET Transwell inserts (CellTreat, Cat no: 230635) were coated with 50 ug/ml collagen I (Gibco, Cat no: A10483-01). Colon organoids cultured 7 days after the previous passage were collected. After centrifugation at 1000G for 5 min at 4°C, The Matrigel/organoid pellet was dissolved with Cell Recovery solution on ice for 45-60 min. After centrifugation at 1000G for 5 min at 4°C, the organoid pellet was resuspended with pre-warmed TrypLE Express (Gibcon, Cat no: 12604-013) and incubated for 5 min at 37°C. Then, the organoids were mechanically dissociated into single cells. Washing media containing advanced DMEM/F12 (Gibco, Cat no: 12634-010), 9% HI-FBS (Cytiva HyClone, Cat no: SH30910.03), 2mM Glutamax (Gibco, Cat no: 35050-061), and 100 unit/mL Penicillin-Streptomycin (PS) (Gibco, Cat no: 15-140-122) was added to neutralize the enzyme and centrifuged at 300G for 5 min at 4°C. The organoid pellet was resuspended in the organoid culture media without nicotinamide as a colon monolayer seeding media. 1 x 10^5^ cells per well were seeded onto collagen I-coated Transwell inserts. 600μl of seeding media was added onto the basolateral side of each transwell, and monolayers were incubated at 37°C. Their growth and confluence were evaluated every day. After 3-4 days of seeding, the first media change was done, and transepithelial electrical resistance (TEER) measurements were performed using the EndOhm-6 chamber with an EVOM2 epithelial voltohmmeter (World Precision Instruments). TEER values were evaluated daily until the day of the experiment to confirm the monolayer’s confluency and establishment of the monolayer culture.

#### Cell sorting of tissue-resident immune poopulations

Gut-IELs, Gut-LP cells, and Liver immune cells were stained with LIVE/DEAD fixable green, fluorescent reactive dye (ThermoFisher, Cat no: L34970) according to the manufacturer’s protocol after tissue processing. In the meantime, the antibody mixture was prepared in BD Horizon Brilliant Stain Buffer ((BD Bioscience, Cat no: 563794). Then, cells were further stained with BV510-conjugated mouse anti-human CD45 (1:20) (BD Bioscience, Cat no: 563204) and PE-conjugated mouse anti-human CD3 (1:10) (BD Bioscience, Cat no: 555333) followed by incubation with human BD Fc block (1:100) (BD Bioscience, Cat no: 564220). After staining at 4°C for 20 min, cells were washed with FACs buffer containing 1%BSA in PBS and centrifuged at 300G for 10 min at 4°C. Cells were resuspended in FACs buffer, and CD45+ cells from IELs, LPs, and Liver-immune cells were separately sorted by fluoresce-activated cell sorter (Cytoflex SRT, Beckman Coulter).

#### Isolation of gut tissue-resident immune cells from colon tissue

The co-cultured system with gut tissue-resident immune cells was established for the Gut-MPS-MOTIVE compartment. To do this, intraepithelial lymphocytes (IELs) were isolated from the epithelial layer of the colon tissue, and lamina propria immune cells were isolated from lamina propria (LP) of the colon tissue.

##### Isolation of IELs

Cryopreserved colon tissue pieces were thawed swiftly at 37°C, washed with the rinse/thawing buffer, and transferred to a 50 ml tube containing the rinse/thawing buffer. Then, it was centrifuged at 400xG for 10 minutes. The supernatant was discarded, and the pellet was resuspended with 15 ml of prewarmed predigesting buffer by inversion. Then, it was incubated horizontally for 20 min at 37°C under continuous rotation (140 RPM) using an incubating shaker (Thermo Scientific, MaxQ 420HP). After that, the tissue pieces waited to settle to the bottom of the 50-ml tube, and then the supernatant containing the stripped epithelium was transferred into a new 50-ml conical tube placed on ice using a pre-wetted 25 ml serological pipette. Another 15ml of fresh prewarmed predigesting buffer was added to the colon tissue pieces and incubated horizontally for 20 min at 37°C under continuous rotation (140 RPM) using an incubating shaker (Thermo Scientific, MaxQ 420HP). After the second round of epithelial stripping, the tube was mixed gently for 10 seconds using a vortex and then waited to be settled at the bottom of the 50-ml conical tube. The second supernatant containing the stripped epithelium and IELs was collected and combined with the first round into a 50 ml conical tube placed on ice using a pre-wetted 25 ml serological pipette. The 70μm cell strainer was washed with 5-10ml of rinse buffer, and the combined supernatant was passed through a 70μm cell strainer placed on a new 50-ml conical tube. 15 ml of ice-cold Rinse buffer was added to the tissue pellet, then centrifuged at 400xG for 10 minutes. The supernatant was transferred to a tube with a 70μm cell strainer and combined with previous supernatants. This combined cell suspension containing stripped epithelium and IELs was centrifuged at 500g for 10 minutes. The cell pellet was resuspended with rinse buffer. Total cell number and cell viability were determined with an automated cell counter (Denovix, CellDrop FL) using trypan blue staining.

##### Isolation of immune cells from the lamina propria of colon tissue

The remaining tissue pieces were transferred to gentleMACS^TM^ C tubes (Milteny Biotec, Cat no: 130-093-237) and washed with rinse buffer. Then, they were centrifuged at 500G for 5 min. The digestion buffer was prepared as follows: 5 mg/ml Collagenase D (Roche; Cat no: 11088858001), 0.5U/ml Dispase (Gibco; Cat no: 17105-041), 30 μg/ml DNAse I (StemCell Technologies, Cat no: 07470), 10 mM HEPES (Gibco; Cat no: 15630-080), 1% PSA (Lonza, Cat no: 17-745E) in HBSS with calcium and magnesium (Cytiva, Cat no: SH30030.01). 2.5 ml of prewarmed digestion buffer was added to tubes and incubated for 30 min at 37°C under continuous rotation (140 G) using an incubating shaker. Then, mix for 10 using a vortexer. The C tubes with the digested pieces were placed into the gentleMACS^TM^ Dissociator, and the program ‘‘m_intestine_01’’ was used twice. The tubes were briefly spun, and samples were aspirated using a 5-ml syringe and a blunt 20G needle three times. The cell suspension was passed through a 70μm cell strainer and washed with rinse buffer. The same digestion step was repeated by adding another 2.5 ml pre-warmed digestion buffer to the remaining tissue pieces and incubating for 30 min at 37°C under continuous rotation (140 G) using an incubating shaker. Then, mix for 10 using a vortexer. The C tubes with the digested pieces were placed into the gentleMACS^TM^ Dissociator, and the program ‘‘m_intestine_01’’ was used twice. The tubes were briefly spun, and samples were aspirated using a 5-ml syringe and a blunt 20G needle three times. The cell suspension was passed through a 70μm cell strainer and washed with rinse buffer. All collected lamina propria cells were combined, and total cell number/viability was determined with an automated cell counter (Denovix, CellDrop FL) using trypan blue staining.

The IELs and LP cell suspensions were centrifuged at 500G for 15 min separately. The cell pellets were resuspended in a DNAse solution prepared with 100 μg/ml DNAse I (StemCell Technologies, Cat no: 07470) in RPMI-1640 and incubated at room temperature (RT) for 15 min. An equal volume of rinse buffer was added, and the suspensions were centrifuged at 300G for 10 min.

#### Gut tissue-resident immune cells culture

FACs sorted Gut-LPs (30.000-50.000 cells/well) from both donors were seeded onto 12-well 0.4 μm PET Transwell inserts (CellTreat, Cat no: 230621) and cultured in OpTmizer™ CTS™ T-Cell Expansion SFM (Gibco, Cat no: A1048501) supplemented with 2mM Glutamax (Gibco, Cat no: 35050-061), 100 units/ml Penicillin-Streptomycin (Gibco, Cat no: 15-140-122), IL-2 (50UI) (R&D Systems, Cat No: 202-IL/CF), and ImmunoCult Human CD3/CD28 T cell activator (Stemcell Technologies, Cat no: 10991) a day before starting co-culture experiment. FACs sorted Gut-IELs were cultured overnight in a T25 cell culture flask with the same media composition. On the day of the experiment, IELs were collected and centrifuged at 300G for 10 min. IELs were resuspended in the apical media consisting of Advanced DMEM/F12 (Gibco, Cat no: 12634-010), HEPES, 2mM Glutamax (Gibco, Cat no: 35050-061), 1x penicillin-streptomycin-amphotericin B (PSA) (Lonza, Cat no: 17-745E), and IL-2 (50UI) (R&D Systems, Cat No: 202-IL/CF). 20.000-40.000 IELs per transwell were divided into each apical side of transwells for both donors.

#### Hepatocyte sandwich culture

Primary hepatocyte sandwich cultures were initiated based on the protocol described previously ^47, 48^. All the reagents were kept on ice. The Matrigel® (Corning, Cat no: 356231) was dissolved in PBS in a 1:100 ratio by vortexing. 24-well 0.4 μm PET Transwell inserts (CellTreat, Cat no: 230635) were coated with 100ul of dissolved Matrigel and incubated for two hours at 37 °C. Donor-matched primary cryopreserved human hepatocytes were purchased from LifeNet Health, who also provided us with the complete donor tissues. Hepatocytes were thawed rapidly in the water bath and centrifuged in hepatocyte thawing media (Lonza, Cat. no.: MCHT50) at 100 G for 5 minutes. The supernatant was discarded, and the pellet was resuspended into the required volume of hepatocyte plating media (Lonza, Cat. No.:MP250). The cell count was then performed using trypan blue on the cell counter. After Matrigel coating was completed, excess solution was removed from each insert, and 800,000 cells seeded in 200ul of plating media. 600ul of plating were also added to the basolateral sides. The seeded transwell inserts were then incubated for 12 hours at 37C in the 5% CO2-supplied incubator. After the incubation, the seeding media were removed from the apical and basolateral area and hepatocyte monolayer overlaid with 2mg/ml rat collagen I (Advanced Biomatrix, Cat no:5153) for 3D coating. Incubate the inserts at 5% CO2 for 10-15 mins to solidify the collagen. The hepatocytes sandwich assembly was maintained in the William’s E maintenance media (Gibco, Cat no: A1217601), with 11,11mM glucose and 1 x HCM^TM^ Singlequots^TM^ Growth Supplement Pack (Lonza, Cat no: CC-4182) containing ascorbic acid, transferrin, HEGF, hydrocortisone, BSA (fatty acid-free), 800 pM insulin for 3 days prior to interaction studies.

On the day of interaction studies, the hepatocyte sandwich-cultured with maintenance media was switched to serum-free common media containing William’s E medium (Thermo Fisher, cat. no. A1217601) with 11.11 mM glucose, 1 x HCM^TM^ Singlequots^TM^ Growth Supplement Pack (Lonza, Cat no: CC-4182) (1x ascorbic acid, 1x transferrin, 1x HEGF, 1x hydrocortisone, 1x BSA (fatty acid-free), 800 pM insulin), 1 % Pen/Strep, 50 UI/ml IL-2 (R&D Systems, cat. no. 202-IL) and ImmunoCult Human CD3/CD28 T cell activator (Stemcell Technologies, Cat no: 10991).

#### Isolation of liver-tissue resident immune cells from whole liver tissue

The liver tissue was washed with RPMI buffer to remove excess blood upon receiving the liver tissue. The liver wedge was cut into fine pieces of approximately 3-5mm using a scalpel and sharp scissors at room temperature. In parallel, cryovials were prepared to distribute and store the tissue pieces in 10% DMSO (Tocris, Cat. no: 67685) + 90% heat inactivated FBS (Cytiva, Cat. no: SH30071.03HI) freezing media. Each vial was labeled properly, stored at −80 °C for 24 hours, and then transferred to −190 °C for further use. Before the day of isolating the immune cell from cryopreserved liver tissue, different buffers were prepared, such as Rinse buffer (RPMI-1640 (Gibco, cat. no: 11875093) + 1% PSA (Lonza, Cat. no: 17745E) + 5% FBS), Digestion buffer (0.5mg/ml Collagenase D () + DNase 30ug/ml + 10mM HEPES (Gibco, 15630-080) + PSA 1% + HBSS (Ca+ Mg+) (Cytiva, Cat. no: SH30030.02)) and EGTA buffer (10mM EGTA (Invitrogen, Cat. no: AM9260G) + HBSS (Ca-, Mg-)) and stored them in 4°C. The next day, all the buffers were placed in a 37 °C water bath, and cryovials having liver tissue pieces removed from −196 °C, were kept on ice and immediately thawed in a 37 °C water bath for 2 minutes. Thawed tissues were transferred to 30 ml of warm RPMI-1640 media and centrifuged for 10 minutes at 300 × g at room temperature (RT). The supernatant was discarded, and the pellet was resuspended into the 20 ml EGTA buffer. The sample was shaken for 20 minutes at 37 °C in the shaker incubator. The samples were added 20ml of Rinse buffer to a 40ml total volume. The sample was centrifuged for 10 minutes at 300xg RT. The pellet was resuspended in a 2.5 ml digestion buffer and shaken for 20 minutes at 37 °C in the shaker incubator. Thereafter, digested samples were transferred to Miltenyi C-tubes to homogenize the tissue using the GentleMACS tissue dissociator preset program E.01. The homogenized samples were transferred to a 50 mL tube using a 100 μm pore size filter. We rinsed the filter with rinse buffer to a total of 40ml. The samples were then centrifuged for 10 minutes at 800G at 4C. The supernatant was decanted, the pellet was resuspended in 30 ml 33% Percoll solution (RPMI67% 33% Percoll) (Cytiva, Cat no: 17544501), and the samples were spun at 1000g for 20 minutes at RT with acceleration and brake set to 1 and 0, respectively. The supernatant was gently decanted, and the pellet was resuspended in 5ml AKC lysis buffer (Gibco, Cat no: A10492-01) for 1 min and then centrifuged 800xg for 5 min at 4C. The supernatant was gently discarded, and the pellet was resuspended in RPMI solution (RPMI+ 1% PSA+ 10% FBS+ 15mM Glutamax) and again centrifuged at 800xg for 5 minutes at 4°C. The pellet was resuspended in DNAse I solution (StemCell Technologies, Cat no: 07470), incubated for 15 minutes at RT, and centrifuged at 500xg for 10 mins at RT. The supernatant was discarded, and the pellet was resuspended in 1-2ml (based on the density of the pellet) PBS solution. The obtained cells were counted using trypan blue.

##### Liver tissue-resident immune cells (LTRL) culture

FACS sorted liver-resident CD45+ immune cells (20.000-40.000 cells/well) were seeded onto 12-well 0.4 μm PET Transwell inserts (CellTreat, Cat no: 230621) and cultured in OpTmizer™ CTS™ T-Cell Expansion SFM (Gibco, Cat no: A1048501) supplemented with 2mM Glutamax (Gibco, Cat no: 35050-061), 100 units/ml Penicillin-Streptomycin (Gibco, Cat no: 15-140-122), IL-2 (50UI) (R&D Systems, Cat No: 202-IL/CF), and ImmunoCult Human CD3/CD28 T cell activator (Stemcell Technologies, Cat no: 10991) a day before starting co-culture experiment.

### MOTIVE PLATFORM AND INTERACTION STUDIES

#### Fabrication and Assembly

The developed MPS system has two components: 1) Two compartment Multiorgan tissue interaction vessel (MOTIVE-2) and 2) the MOTIVE platform, supporting MOTIVE-2 perfusion. The body of MOTIVE and supporting platform were designed in CAD and printed on the in-house Stratasys Polyjet-J750 3D printer. Biocompatible Digital Acrylonitrile butadiene styrene (ABS) Plus (Stratasys, Cat. no: OBJ:03721) material was used to print the MOTIVE and the platform. The dimensions of vessel and platform were estimated to (3.09’ X 1.33’ X 1.10’) and (7.59’ X 7.59’ X 1.80’), respectively. The vessel has two 0.39’ apart compartments (0.8’ diameter, 0.86 depth) designed to fit 12-well membrane inserts. Flow of media between the compartment in the vessels was achieved through 3.3V minipumps (0.57’ X 0.50’ X 0.69’) supplied by Takasago fluidics system (Cat. no: 2006097296) and the pharma grade silicone tubing (0.063’ X 0.0125’ X 50’) (Thermo Scientific, Cat. no: 8600-0020). The connections between the pumps and tubing were made through male and female barbed Luer locks (0.10’ X 0.15’) which is linked with the inlet and outlet (0.14’ dia) of MOTIVE. The supporting perfusion platform has three installed 3.3V minipumps controlled by a custom-fabricated controller and paper ink display (RSEN s.p., Slovenia). The pumps are connected in series with a 14.4V rechargeable battery. The single platform can support the operation of three MOTIVEs with tested flowrate of 2.60 ml/min and 0.05 MPa discharge air pressure.Prior to the interaction experiment each MOTIVE-2 was plasma sterilized for 4 hrs, coated with Biofloat coat for 3mins (faCellitate, Cat no: F202005) and primed with glucose free WEM media for 12hrs. The colon and liver immune co-cultures in transwell were then transferred to the MOTIVE-2 into the designated compartment, covered with a lid, attached to the peristaltic pumps on the MOTIVE platform, and incubated at 37C 5% CO_2_-supplied incubator for two days of interaction. After the experiment, the samples were collected for further analysis.

#### Establishing a co-cultured Gut MPS system with tissue-resident immune cells using MOTIVE

Colon epithelial monolayers with TEER higher than 1000 ohm/cm^2^ were selected for interaction and isolation studies. The seeding media was switched to the serum-free apical media containing Advanced DMEM/F12 (Gibco, Cat no: 12634-010), HEPES, 2mM Glutamax (Gibco, Cat no: 35050-061), 1x penicillin-streptomycin-amphotericin B (PSA) (Lonza, Cat no: 17-745E), and IL-2 (50UI) (R&D Systems, Cat No: 202-IL/CF). The colon epithelial monolayers in the Gut-MPS compartment were maintained in this apical media during the time course of the experiment. Gut-IELs were added to the apical side of the 24-well transwell inserts that have colon epithelium monolayer on the transwell membrane. These 24-well transwell inserts were aimed to be placed in 12-well transwell inserts that consist of LP cell suspensions separated by a 0.7’ O-ring and hung into the media reservoir to establish the co-cultured MPS system of the colon epithelium with its own gut tissue-resident immune cells. The basolateral gut compartment, as well as the basal liver compartment, were fluidically connected to systemic circulation on the MOTIVE platform using serum-free common media containing William’s E medium (Thermo Fisher, Cat no: A1217601) with 11.11 mM glucose, 1 x HCM^TM^ Singlequots^TM^ Growth Supplement Pack (Lonza, Cat no: CC-4182) containing 1x ascorbic acid, 1x transferrin, 1x HEGF, 1x hydrocortisone, 1x BSA (fatty acid-free), 800 pM insulin, 1 % Pen/Strep. 50 UI/ml IL-2 (R&D Systems, cat. no. 202-IL) and ImmunoCult Human CD3/CD28 T cell activator (Stemcell Technologies, Cat no: 10991) were also added in the common media.

#### Establishing a co-cultured Liver MPS system with tissue-resident immune cells using MOTIVE

The same donor’s human liver primary hepatocytes were purchased from LifeNet Health LifeSciences. 3-4 days before interaction and isolation experiments, the hepatocytes were seeded onto 24-well transwell inserts using a sandwich method as described above. A day before the experiment, liver tissue-resident immune cells were isolated from liver tissue and seeded onto 12-well transwell inserts. 24-well transwell inserts containing a sandwich culture of the human liver primary hepatocytes were stacked in 12-well transwell inserts that consist of liver tissue-resident cell suspensions separated by a 0.7’ O-ring and hung into the media reservoir to establish the co-cultured MPS system of the liver hepatocytes with its own liver tissue-resident immune cells. The basolateral liver compartment, as well as the basal gut compartment, were fluidically connected to systemic circulation in MOTIVE-2 vessels.

#### Operation of the MOTIVE platform and conducting MPS interaction studies

On the day of the experiment, all tubings were primed with serum-free common media. While pumping the common media, the pump function and tubing connections from the pump to each dry compartment were visually confirmed. 10 ml serum-free common media was added to each MOTIVE platform. Common media was prepared with William’s E medium (GIBCO, cat. no. A1217601) containing 11.11 mM glucose, 1 x HCM^TM^ Singlequots^TM^ Growth Supplement Pack (Lonza, Cat no: CC-4182), 1x ascorbic acid, 1x transferrin, 1x HEGF, 1x hydrocortisone, 1x BSA (fatty acid-free), 800 pM insulin, 1 % Pen/Strep and 50 UI/ml IL-2 (R&D Systems, cat. no. 202-IL) and the ImmunoCult Human CD3/CD28 T cell activator. Gut MPS and Liver MPS compartments were established in the double-transwells setting. 12-well transwell inserts containing LP cell suspensions (400μl) were initially placed in the MOTIVE platform for the Gut MPS compartment. Then, 24-well transwell inserts with a colon epithelium monolayer and Gut-IEL cell suspension (200μl) were inserted in the 12-well inserts to assemble the Gut-MPS on the MOTIVE-2 vessel. The same approach was applied to Liver MPSs as follows: 12-well transwell inserts containing liver-immune cell suspensions (400μl) were initially placed in the Liver-MPS compartment of the MOTIV-2. Then, 24-well inserts with a human hepatocyte sandwich culture (200μl) were inserted.

##### Immune activation challenge

Lipopolysaccharide (LPS) from *Escherichia coli (E. coli)*, Poly (I: C), and 5-(2-oxopropylideneamino)-6-D-ribitylaminouracil (5-OPRU) as immune activators to discern donor specific immune responses. Apical gut compartments in the interactions received either1μg/ml of LPS (Novus biologicals, Cat no: NBP2-25295), 1μg/ml of Poly (I: C) (Tocris, Cat no: 4287), or 8000nM of 5-OP-RU (GVK Biosciences Pvt. Ltd.) Each inflammatory activator was added to the apical side of the 24-well transwell inserts containing the colon epithelium monolayer and Gut-IELs. Then, they were further placed into 12-well transwell inserts containing Gut-LP cell suspensions. Interaction studies with the immune activators or controls without were performed in fluidic communication with the liver/immune MPS in MOTIVE-2 vessels for 2 days.

The morphology of each co-cultured colon epithelium and hepatocytes post-treatment was analyzed with an inverted microscope (Nikon Eclipse Ti-S). TEER was measured before starting interaction and 2 days after interaction experiments to monitor the integrity of the gut barrier. At the end of the interaction studies on day 2, media from the apical gut and liver compartments was collected. Common media, which is in the systemic circulation reservoir, was also collected. Gut-IEL, Gut-LP, and Liver tissue-resident immune cells were collected with the media in a low retention sterile microcentrifuge tube (Thermo Scientific, Cat no: 3451) and spun down at 500xg for 5 minutes. The supernatant was collected in low protein-binding tubes (Thermo Scientific, Cat no: 88379), and 0.5% BSA was added. Samples were transferred to a −80°C freezer for further analysis of cytokine/chemokine and albumin assays. The pellets, including tissue-resident immune cells, were also stored in low-retention sterile tubes and transferred to a −80°C freezer for RNA isolation. The colon monolayers and immune cells on the transwell membranes and hepatocytes were collected in a low retention sterile microcentrifuge tube (Thermo Scientific, Cat no: 3451) within either TRIZOL reagent or lysis buffer and transferred to a −80°C freezer for further RNA isolation. Each condition was performed as two 2 biological replicates per individual donor.

#### Isolation control studies

Gut-Isolation (ISO) and liver-Isolation (ISO) studies were conducted with 12-well plates without using the MOTIVE platform but with the same double-transwell setting. These two transwells were assembled onto 12-well plates on the same day of the interaction study. Isolation experimentation was carried out with different settings as follows: i) co-cultured with tissue-resident immune cells and no treatment (ISO-1), ii) co-cultured without tissue-resident immune cells (ISO-2), iii) tissue only with Poly (I: C), iv) tissue only with LPS, v) tissue only with 5-OPRU.

##### Gut-Isolation (ISO) study

The colon monolayer seeding media was removed upon selection based on the TEER measurement. Gut-IELs were resuspended within gut apical media consisting of Advanced DMEM/F12 (Gibco, Cat no: 12634-010), HEPES, 2mM Glutamax (Gibco, Cat no: 35050-061), 1x penicillin-streptomycin-amphotericin B (PSA) (Lonza, Cat no: 17-745E), and IL-2 (50UI) (R&D Systems, Cat No: 202-IL/CF). Then, the same number of Gut-IELs as the interaction study was added to the apical side of the 24-well transwell inserts with colon epithelium monolayer on the transwell membrane. These 24-well transwell inserts were placed in 12-well transwell inserts with the same number of LP cells as the interaction study to establish the co-cultured Gut-Isolation study co-cultured with tissue-resident immune cells (ISO-1). The basolateral side of the 12-well plates was filled with 1.5 ml of serum-free common media supplemented with 50 UI/ml IL-2 (R&D Systems, cat. no. 202-IL) and ImmunoCult Human CD3/CD28 T cell activator, which is the same media used in the gut compartment of interaction study. For the Gut-Isolation without co-culture of the immune cells (ISO-2), colon epithelium monolayers were placed in the 12-well transwell inserts containing only 400μl serum-free common media with 50 UI/ml IL-2 (R&D Systems, cat. no. 202-IL) without immune cells. The basolateral side of the 12-well plates was filled with 1.5 ml of serum-free common media supplemented with 50 UI/ml IL-2 (R&D Systems, cat. no. 202-IL).

##### Liver-Isolation (ISO) study

On the same day as the interaction study, primary hepatocytes cultured with the sandwich method onto a 24-well transwell were also selected for isolation studies. For the cocultured Liver-Isolation study with tissue-resident immune cells (ISO-1), the apical media was changed to serum-free common media supplemented with 50 UI/ml IL-2 (R&D Systems, cat. no. 202-IL) and ImmunoCult Human CD3/CD28 T cell activator, which is the same media used in the interaction study. These 24-well transwell inserts with sandwich-cultured hepatocytes were placed in 12-well transwell inserts with the same number of liver-tissue resident immune cells (LTLs) as in the interaction study. The basolateral side of the 12-well plates was filled with 1.5 ml of the same media composition within the apical side of the transwell, which is also the same media used in the liver compartment of the interaction study. To assemble Liver-Isolation without co-cultured immune cells (ISO-2), the sandwich-method cultured primary hepatocytes apical media was switched to 200μl of serum-free common media with 50 UI/ml IL-2 (R&D Systems, cat. no. 202-IL). Then, they were put in the 12-well transwell inserts containing only 400μl of serum-free common media with 50 UI/ml IL-2 (R&D Systems, cat. no. 202-IL). The basolateral side of the 12-well plates was filled with 1.5 ml of serum-free common media supplemented with 50 UI/ml IL-2 (R&D Systems, cat. no. 202-IL).

### ANALYTICAL PROCEDURES

#### Multiplex cytokine and chemokine assays

Cytokine and chemokine levels were quantified post 2 days of exposure to the inflammatory conditions using Human Custom ProcartaPlex 27-plex (Invitrogen, Cat no: PPX-27-MX2XAPJ) and Human Custom ProcartaPlex 22-plex (Invitrogen, Cat no: PPX-22-MXT2CGF). The samples were analyzed using the Luminex™ 200™ Instrument System, and the multiplexed cytokine/chemokines data was collected using the xPONENT FLEXMAP 3D software (Luminex Corporation, Austin, TX, USA). The concentration of each analyte was calculated from a standard curve, which was generated by fitting a 5-parameter logistic regression of mean fluorescence on known concentrations of each analyte. The TGF-β 1,2,3 production level was separately quantified using MILLIPLEX TGF-β 1,2,3 magnetic bead kit (Millipore, Cat no: TGFBMAG-64K-03) analyzed with Luminex™ 200™ Instrument System.

#### ELISA albumin assay

To assess the performance of hepatocytes during interaction and isolation, samples were collected from various conditions. The samples were analyzed using Human albumin ELISA kit (Raybiotech, Cat. no: ELH-Albumin-1). Assays were performed as per the manufacture’s guidelines.

#### Immunofluorescence assay and Imaging

Transwell membranes with colon epithelial monolayers were washed three times with 1x DPBS (Gibco, Cat no:14190-144). Then, cells were fixed with 4% paraformaldehyde (ChemCruz, Cat no:sc-281692) for 20 min at room temperature (RT) and washed with 1x PBS. Permeabilization was done using 0.1% Triton X-100 (Thermo Scientific, Cat no: A16046.AE) in 1x DPBS and incubated at RT for 15 min. After removing Triton X-100, cells were washed with 1x DPBS, and blocking buffer (2% BSA in 1xPBS) was added and incubated at 4°C overnight. Then, cells were stained with primary LGR5 monoclonal antibody (OTI2A2) (1:100) (Invitrogen, Cat no: MA5-25644) for 1-3 hours at room temperature. After washing three times with 1x PBS, cells were stained with Alexa Fluor™ 488 Goat anti-Mouse IgG (H+L), Superclonal™ Recombinant Secondary Antibody (1:1000) (Invitrogen, Cat no: A28175) for 45 min at RT. For the staining of goblet cells and enteroendocrine cells, transwell membranes containing colon epithelial monolayers were separately stained with Mucin 2/MUC2 Antibody (F-2) conjugated with Alexa Fluor® 647 (1:100) (Santa Cruz, Cat no: sc-515032 AF647). Chr-a Antibody (C-12) conjugated with FITC (1:100) (Santa Cruz, Cat no: sc-393941 FITC). Then, concurrently stained with either Actin-Red (Invitrogen, Cat no: R37112) or Actin-Green (Invitrogen, Cat no: R37110) and NucBlue™ Live ReadyProbes™ Reagent (Hoechst 33342) (Invitrogen, Cat no: R37605). Transwell membranes were cut and mounted on glass slides with ProLong™ Gold Antifade reagent (Invitrogen, Cat no: P36934). Microscopy images from both donor samples were taken with the Leica Thunder at 10x and 20x magnification.

#### RNA isolation, cDNA library preparation, and bulk RNA-sequencing

Colon monolayer epithelial cells with Gut-IEL and Gut-LP from Gut-MPSs and sandwich-method cultured hepatocytes and liver-immune cells from Liver-MPSs were collected. mRNA was extracted using the RNeasy Micro kit (Qiagen, Cat no: 74004) for donor 1 and 2 colon epithelial samples and donor 1 hepatocyte samples. For donor-2 hepatocyte samples and Gut-LP, IEL, and liver-immune cells, mRNA was extracted using TRIZOL reagent (Invitrogen, Cat no: 15596018) according to the manufacturer’s protocol. mRNAs of Donor 1 immune cells were extracted using NucleoSpin RNA XS (Takara, Cat no: 740902.50). Total RNA and RIN number were analyzed using Agilent 2200 TapeStation system with High Sensitivity RNA ScreenTape (Agilent, Cat no:5067-5579), High Sensitivity RNA ScreenTape Ladder (Agilent, Cat no:5067-5581), High Sensitivity RNA ScreenTape Sample Buffer (Agilent, Cat no:5067-5580). Total RNA was quantified using Qubit™ dsDNA Quantification Assay Kits (Invitrogen, Cat no: Q32852) on a Qubit™ Flex Fluorometer instrument (Thermo Fisher).

Libraries were prepared with SMARTer Stranded Total RNA-Seq Kit v2 - Pico Input Mammalian (Takara, Cat no: 634418) along with Indexing primer set HT for Illumina v2 (Takara, Cat no: 634421) and SMARTer® RNA Unique Dual Index Kit-24U (Takara, Cat no: 634451). Then, the resulting libraries were analyzed using a 2200 TapeStation system (Agilent, G2964AA) with High Sensitivity D1000 ScreenTape System (Agilent, Cat no: 5067-5584) and High Sensitivity D1000 Reagents (Agilent, Cat no: 5067-5585) and quantified using qPCR using Kapa Library Quantification Kit (Roche, Cat no: KK4824). Then, pre-made libraries were pooled before sequencing with NovaSeq X Plus PE150 by NovoGene at an average depth of 36 million reads per sample.

#### RNA-seq Data Quality Control and Pre-Processing

Adapters were removed from raw paired-end RNA-seq FASTQ files using Cutadapt (version 4.9) with default parameters. TruSeq adapter sequences were provided as a FASTA file by the manufacturer. Trimmed paired-end reads were aligned to the human reference genome (GRCh38, Ensembl release 112) and quantified using RSEM (v1.3.3) with STAR as the aligner. The reference genome and gene annotation (GTF) files were downloaded from Ensembl and used to build the RSEM reference via rsem-prepare-reference with STAR support. Expression quantification was performed with rsem-calculate-expression using default parameters and STAR as the aligner. RSEM was run in paired-end mode with the −-estimate-rspd, −-append-names, and −-output-genome-bam flags enabled. Gene- and isoform-level expression values were extracted for downstream analyses under the result file column “expected count”.

#### RNA-seq Differentially Expressed Gene Analysis

Differential expression analysis was performed separately for each donor and cell type using DESeq2 (version 1.42.1) with raw gene-level count data. Genes with fewer than 10 total counts across all samples in a given comparison were excluded. For treatment-based comparisons on the full gut-liver-on-a-chip device, DESeq2 datasets were constructed using the design formula ∼ Treatment to evaluate the effects of antigen exposure within each individual cellular compartment. For device comparison studies involving the same meta cell types (excluding the epithelial chamber) in the absence of antigen treatment, DESeq2 datasets were constructed using the design formula ∼ Device, where the device condition captured four distinct configurations: (1) isolated meta cell type cultures, (2) tissue-immune cell co-cultures without gut-liver interconnection, (3) gut-liver-connected devices without immune cells, and (4) gut-liver-connected devices including immune components. For comparisons involving the epithelial chamber, a separate DESeq2 dataset was created that included epithelial cells and intraepithelial CD45⁺ cells under various configurations: epithelial cells alone, intraepithelial CD45⁺ cells alone, epithelial and intraepithelial CD45⁺ co-cultures, epithelial cells in gut-liver-connected devices without immune components, and epithelial–intraepithelial CD45⁺ co-cultures in the fully connected model. Genes with an adjusted p-value < 0.05 and an absolute log2 fold-change > 1 were considered significantly differentially expressed.

#### Gene Set Enrichment Analysis

Enrichment analysis was performed on upregulated and downregulated genes using ontology-wide GSEA was conducted using the gseGO function from clusterProfiler to explore enrichment across all major GO domains (biological process, molecular function, and cellular component). Pathways and gene sets with a Benjamini-Hochberg adjusted p-value < 0.05 were considered significantly enriched. GSEA Pathway enrichment results were expressed as normalized enrichment scores (NES), p-values, and gene set sizes.

Pathway enrichment results (normalized enrichment scores (NES), p-values, and gene set sizes) were visualized using a custom Python script. Briefly, the data were read into a Pandas (v1.4) DataFrame and processed with NumPy (v1.22). P-values were transformed to –log₁₀(p-value) for color mapping.

Bubble plots were generated with Matplotlib (v3.5). The x-axis shows NES, the y-axis lists pathway names sorted in ascending order of reversed NES, and each bubble’s area is proportional to the gene set size (setSize × 0.5). Bubble color encodes –log₁₀(p-value) using the Viridis colormap. To produce a compact layout suitable for manuscript figures, the canvas was set to 12 × 10 inches, and vertical spacing between pathway labels was fixed at 0.02 units. Axis labels, tick labels, and titles were rendered at 20 pt and 26 pt font, respectively; annotations (the numeric NES values) were added above each bubble with an upward offset of 10 points and horizontal offsets of +5 or –20 points (depending on proximity to the plot edges), on a semi-transparent white background to ensure legibility. Figures were exported at 300 dpi.

#### Single-cell RNA-seq

The Chromium Next GEM single-cell 5′ Reagent kit v2 (10x Genomics) was used for single-cell RNA sequencing (sc-RNA-seq). Whole tissue and sorted CD45+CD3+ T cells from the gut, liver were loaded onto different lanes of the Next GEM Chromium Controller (10x Genomics) for encapsulation. Single-cell libraries were generated using the manufacturer’s protocols and sequenced by the Johns Hopkins All Children’s Shared Resources Core.

##### scRNA-seq Data Quality Control and Pre-Processing

The scRNA-seq workflow was adapted from our previous work. Seurat^59^ and Scanpy^60^ were used for downstream scRNA-seq data analysis. For quality control, low-quality cells were dropped based on their extremely low UMI counts (<1000/1600/1200 for Donor 1/2/3), high mitochondrial gene counts (>20%), and a small number of uniquely expressed genes (<200) to filter out empty droplet and dead cells. Python package doubletdetection^61^ was used to remove droplets. Scanpy function score_genes_cell_cycle was used to assign each quality-controlled cell with a cell cycle score based on the expression of cell cycle phase genes defined by Triosh et al^62^. Raw counts of quality-controlled cells, along with their cell cycle score and mitochondrial fraction, were input into R for normalization and donor integration. We performed SCTransform for each donor, each organ individually and then integrated different donors together. For our study, cell’s G2/M phase score, S phase score, and mitochondrial percentage were also included in the regression formula to remove the confounding effect of these variables on the gene expression when performing the SCTransform. Regressed gene expression calculated from the regularized generalized linear model coefficients was referred as “SCTransform-corrected counts,” and log-transformed SCTransform-corrected counts were used for later visualization. The Pearson residual of a cell’s observed gene expression to the SCTransform-corrected counts was used for integration and later performing principal component analysis (PCA, 50 pcs kept). Data integration was done by using Seurat, using both donors and one additional donor datasets^5^ to enhance clustering accuracy and mitigate donor-specific effects. While the final organ-on-chip model utilized cells only from donors 1 and 2, we extracted and analyzed their contributions specifically from the integrated dataset for model-relevant characterization. The top 3000 genes that were the most variable across all donors’ organs’ data were selected to define the cell-anchors-finding space and then perform the integration based on canonical correlation analysis. Pearson residuals calculated by the SCTransform were corrected according to the cell anchors. After PCA, Leiden clustering was performed based on the computed neighborhood graphs (50 pcs, size of neighborhood equals 15 cells) to reveal the general subtypes. The initial clustering resolution is set at 0.8. In order to delineate cell subtypes, further subclustering was performed on each subcluster at a resolution range from 0 to 1 (detailed resolution for each step of subclustering was recorded in the published code). UMAP was performed on the neighborhood graph to visualize the clustering (parameters set to scanpy default). These analyses—integration, PCA, UMAP, and cell type annotation—were performed separately for each site (with donor-integrated data).

We classified cells based on their differential expression of the canonical markers summarized from literature: *EPCAM* for intestinal epithelial cells; *ACTA2* for smooth muscle cells; *CD34*, *DCLK1* for fibroblasts; *CCL21* in addition to *CD34* and *DCLK1* for fibroblastic reticular cells which likely in the T cell zone of the mucosal associated lymphoid tissue to attract CCR7+ T cells; *SOX10*, *S100B*, *BMP7* for growth factor-secreting enteric glial cells; *NRG1*, *BMP7*, *WNT5A*, *PDGFRA*, *PDGFRB* for telocytes; *CD34*, *PECAM1*, and *MADCAM1* for endothelial cells; *CD68*, *CD86*, *S100B* for dendritic cells; *CD68*, *CD86*, *CD14*, *MRC1*, *ITGAM*, *FCGR3A* for macrophages; *CD14*, *CD68* but not the aforementioned other macrophages/dendritic cell markers for monocytes; *CD19* for B cells; negative for *CD19* and positive for *IGHA1*, which signifies the class switching of the B cells, for plasma cells; *KIT* for innate lymphoid cells; GATA2, KIT, CD34 for mas progenitors; *TRDC* but not all 4 CD3 positive for natural killer cells; all 4 CD3s (*CD3D*, *CD3E*, *CD3G*, *CD274*) for T cells. For liver-specific cells, we referred to Sonya A. MacParland et al.’s research^3^: CA1 and HBB for erthyroid; RAMP3, LIFR for portal endothelial cells; CD34, PECAM1, MGP, CLEC14A for liver sinusoidal endothelial cells; ACTA2, COL1A1 for stellate cells; KRT19, KRT7, SOX9 for cholangiocytes; GSTA2, AKR1C1, ALB for hepatocytes; SOX10, S100B for glial-like cells.

We further increased the granularity of the T cell annotation (Fig. S5). We classified cells as either TCRαβ or TCRγδ based on the expression of the γδ T cell marker *TRDC* (sufficient when only CD3+ T cells are being considered)^63^. TCRαβ cells were delineated into CD4+, CD8αα+, CD8αβ+, and double negative (DN) subsets based on the expression of *CD4*, *CD8A*, and *CD8B*. We defined six major T cell subtypes as: (1) Central memory T cells (TCM) expressing *SELL*, *CCR7*, *TCF7*, *IL7R*, *SP1R1*, and no tissue residencies markers such as integrins *ITGAE* and *ITGA1*, showing their lymph node homing ability, the self-renewal and differentiation potential, long lives, and circulating ability. (2) Effector memory cells (TEM), which feature impaired *SELL* and *CCR7*, have effector marker *KLRG1*, have *S1PR1* or its transcription factor-encoding gene *KLF2* implies the potential to circulate in the blood, and *IL7R* indicates their long life. (3) Tissue-Resident Memory T cells (TRM), expressing high levels of *IL7R*, *ITGAE*, and *ITGA1*, and low levels of molecules associated with tissue egresses, such as *S1PR1* and *CCR7*. As previously reported, CD4+ TRM expressed lower levels of CD49a (ITGA1) compared to CD8+ TRM^64^. (4) Effector T cells, characterized by low *IL7R* expression (indicating a short-lived phenotype), high *KLRG1* expression, and cytokine production. (5) Mucosal-associated invariant T cells (MAIT), which have a unique semi-invariant TCR α chain that uses *TRAV1-2*.

##### scRNA-seq CellPhoneDB cell-cell interaction analysis

Cell–cell communication inference was performed using CellPhoneDB (version 4.1.0) in statistical mode. For each sample, normalized gene expression data and corresponding cell type metadata were used as input. CellPhoneDB was run with 1,000 permutations, using a 30% expression threshold to include only genes expressed in a minimum fraction of cells within a cell type. Significant ligand–receptor interactions were defined using a p-value cutoff of 0.05. To summarize the frequency of detected interactions, an undirected interaction count matrix was generated by aggregating significant interactions for each unique pair of cell types. Cell–cell interaction frequencies were exported in long and wide formats for chord diagram visualization.

##### scRNA-seq Differentially Expressed Gene Analysis

Differentially expressed gene (DEG) analysis was performed between 2 donors’ T cells and macrophages, respectively, in colon and liver using Monocle 3 (version 1.3.1)^65^. A generalized linear model (GLM) was fitted to SCTransform-corrected counts, assuming each gene’s expression in each cell follows a quasi-Poisson distribution, with its mean and variance as functions of cell’s donor origin. Log_2_ fold change was calculated from the log_2_-transformed ratio between the coefficient of the cell identity variable and the intercept (baseline). The *p*-value was computed using the Wald test on the coefficient to determine whether it was significantly different from zero. Benjamini-Hochberg correction was applied for multiple comparison corrections. The exact calculation procedure is described in the Monocle 3 documentation and our previous work^66^. Genes with adjusted *p*-values < 0.05 and an absolute log_2_ fold change > 1 were considered differentially expressed.

##### NicheNet Ligand Inference and Cell Type Ligand Enrichment Calculations

NicheNet^1^ (version 2.2.0) was applied to infer the ligands most likely responsible for differential gene upregulation associated with the donor specificity in colon and liver T cells and macrophages. DEGs with adjusted p-values < 0.05 and log_2_ fold changes greater than 0.5 were used as input for NicheNet. The integrated ligand-receptor pairs, signaling network, and gene regulatory networks (prefix 21122021) were obtained from the package’s repository. Genes expressed in at least 1% of the cells were considered as background. Potential ligands were defined as ligands of NicheNet-documented receptors whose encoding genes were expressed in at least 1% of the cell subsets. The top 15 ligands with an AUROC > 0.5 were reported in the main figure, while the top 30 were used for quantifying their enrichment in neighboring cell types. If fewer than 15 ligands met the AUROC > 0.5 threshold, all qualifying ligands were visualized and included in the enrichment analysis. The full list of ligands is available in Supplementary Table 3. Each donor’s colon and liver profiles were obtained from averaging scRNA-seq expression by cell type. A gene was included in the averaged profile only if it was expressed in at least 1% of a cell type. For AUC calculation, for each cell type, we iterated down the ranked gene list (ranked by their average expression in the given cell type, descending order) to recover NicheNet-inferred ligands, stopping when encountering a zero-expression gene. The final area under the curve was computed using the auc function from sklearn.metrics.

#### Code availability

Scripts used for transcriptomic analysis can be accessed at https://github.com/Brubaker-Lab/gut-liver-model.

### STATISTICAL ANALYSIS

All experiments were performed at a minimum of two biological replicates and two separate donors.

Concentrations of cytokines and chemokines, TEER and albumin concentrations were ploted and analyzed using GraphPad Prism (Version 10.2.2 (341), GraphPad Software, San Diego, California USA)

Data visualization and calculation of fold changes over the control group for TEER and albumin measurements were performed using GraphPad Prism (Version 10.2.2 (341), GraphPad Software, San Diego, California USA, www.graphpad.com). To determine statistically significant differences between experimental groups for cytokine/chemokine concentrations, TEER values, and albumin levels, a one-way Analysis of Variance (ANOVA) was performed. Post-hoc pairwise comparisons against the designated control group were conducted using Fisher’s Least Significant Difference (LSD) test. Actual p-values are depicted on the graphs. Principal Component Analysis (PCA) and heatmap visualization of cytokine/chemokine data were performed using the web tool ClustVis (https://biit.cs.ut.ee/clustvis/)^43^. For heatmap generation, data were scaled using unit variance scaling and both rows (analytes) and columns (samples) were clustered using correlation distance and average linkage.

Analysis of transcriptomic data from bulk and single-cell RNA sequencing is discussed under their respective methods sections.

### GRAPHICS

Schematics and symbols were created with BioRender (https://BioRender.com).

## Acknowledgements

Single-cell sequencing was performed at the JHU Single Cell and Transcriptomics Core. The study was supported by grants from NIGMS 5R35GM146900 and in part by the Merck Exploratory Science Center Fellowship. R. R. and D.K. B are supported by an award from the Good Ventures Foundation and Open Philanthropy, as well as start-up funds from Case Western Reserve University and University Hospitals.

## Author Contributions

M. U. procured donor tissues, performed experiments, analyzed data, prepared graphs and wrote the manuscript. R. R. analyzed sequencing data, made figures, and wrote the manuscript. J.L. processed samples and analyzed data. M. F. S. performed experiments and operation of the devices, and processed samples for single-cell RNA sequencing. L.R. led the fabrication of 3D printed vessels, M.D. designed and fabricated the device controllers. C.P. processed samples, performed RNA sequencing and analysis, L.L. assisted in study design and data interpretation, D.K.B. assisted in study design and wrote and edited the manuscript. M. T. designed experiments, coordinated the project, and wrote and edited the manuscript.

## Supplementary material

**Figure S1:**
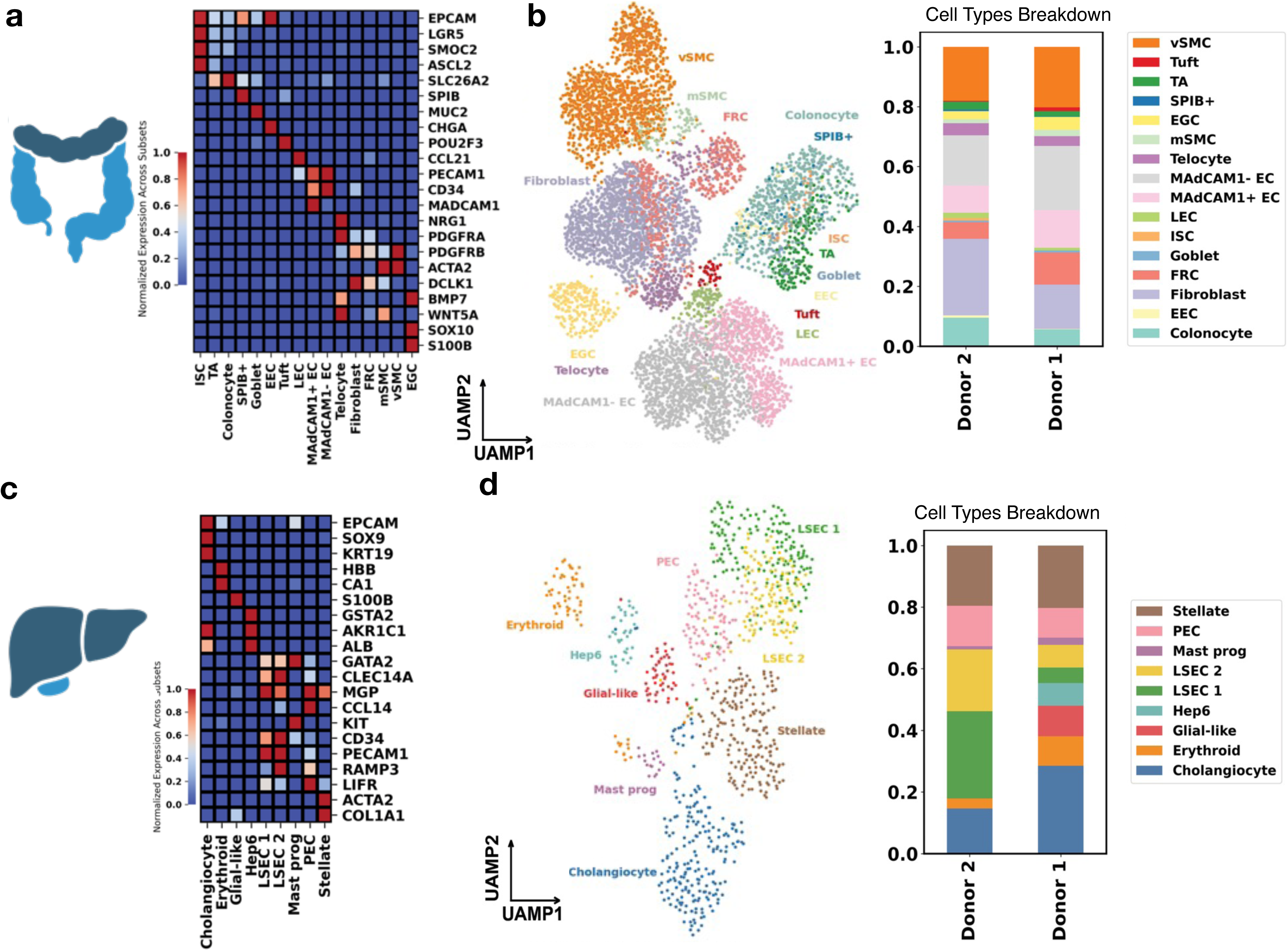
Single-cell characterization of same-donor colon and liver tissue reveals differences in non-immune cell composition. **a,** Marker gene expression in all identified colon non-immune cells. The color of each cell in the heatmap denotes the normalized cell type-averaged expression of each marker. Abbreviations: Colonocyte - colonocytes; EEC - enteroendocrine cells; Fibroblast - fibroblasts; FRC - fibroblastic reticular cells; Goblet - goblet cells; ISC - intestinal stem cells; LEC - lymphatic endothelial cells; MAdCAM1^⁺^ EC - MAdCAM1^⁺^ endothelial cells; MAdCAM1^⁻^ EC - MAdCAM1^⁻^ endothelial cells; Telocyte - telocytes; mSMC - muscularis smooth muscle cells; EGC - enteric glial cells; SPIB^⁺^ - SPIB^⁺^ cells; TA - transit amplifying cells; Tuft - tuft cells; vSMC - vascular smooth muscle cells. **b,** Left: UMAP embeddings of cells collected in the colon of all donors; Right: stacked bar plot shows cell type composition in Donor 1 and Donor 2. **c,** Marker gene expression in all identified liver non-immune cells. The color of each cell in the heatmap denotes the normalized cell type-averaged expression of each marker. Abbrevations: Cholangiocyte - cholangiocytes; Erythroid - erythroid cells; Glial-like - glial-like cells; Hep6 - hepatocytes type 6; LSEC Z1 - liver sinusoidal endothelial cells zone 1; LSEC Z2 - liver sinusoidal endothelial cells zone 2; Mast prog - mast progenitor cells; PEC - portal endothelial cells; Stellate - hepatic stellate cells. **d,** Left: UMAP embeddings of cells collected in the liver of all donors; Right: stacked bar plot shows fractional cell type composition in Donor 1 and Donor 2.

**Figure S2:**
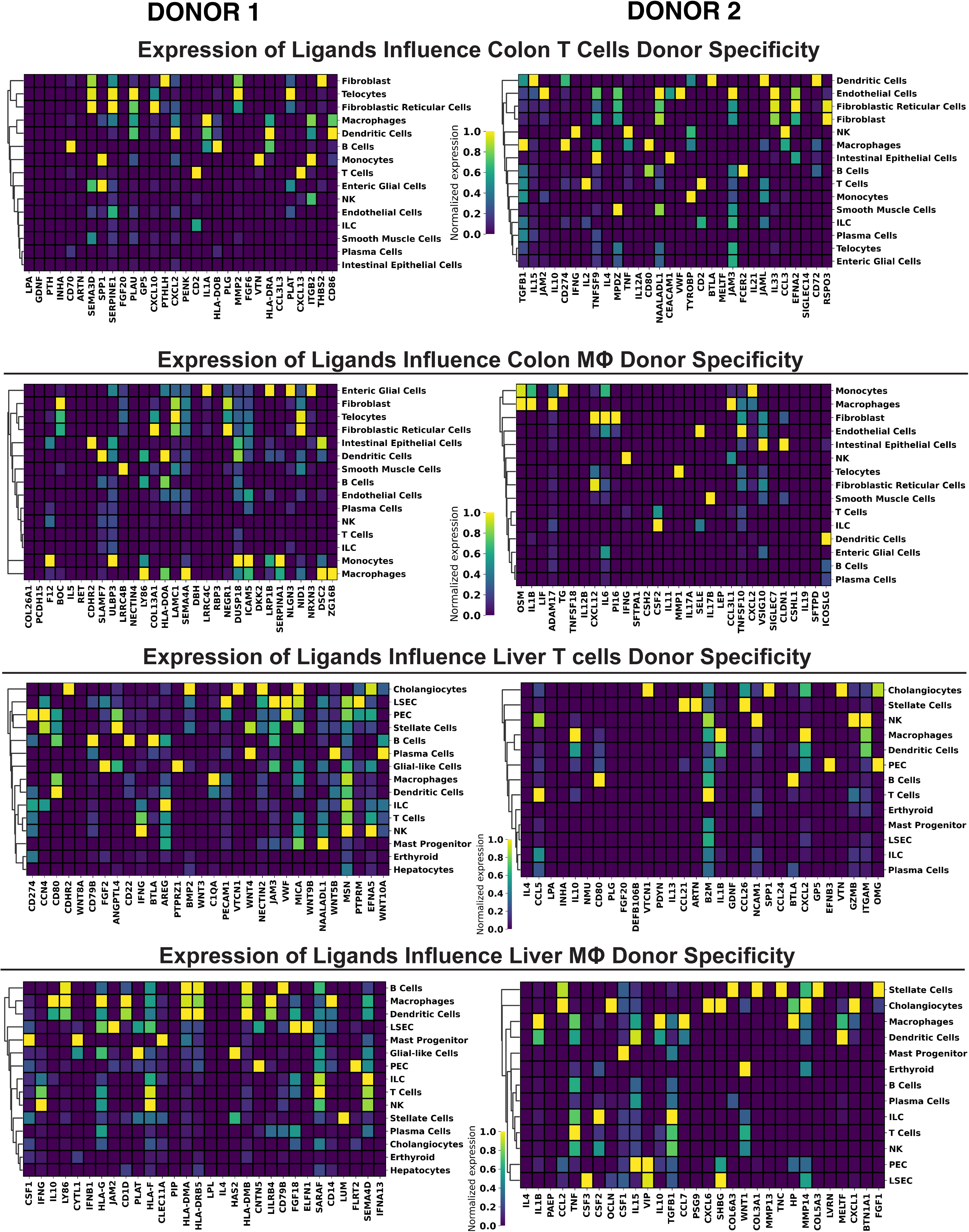
Heatmaps showing the Gene Expression of NicheNet-inferred ligands used for AUC Calculation in each donor’s colon and liver cell types.

**Figure S3:**
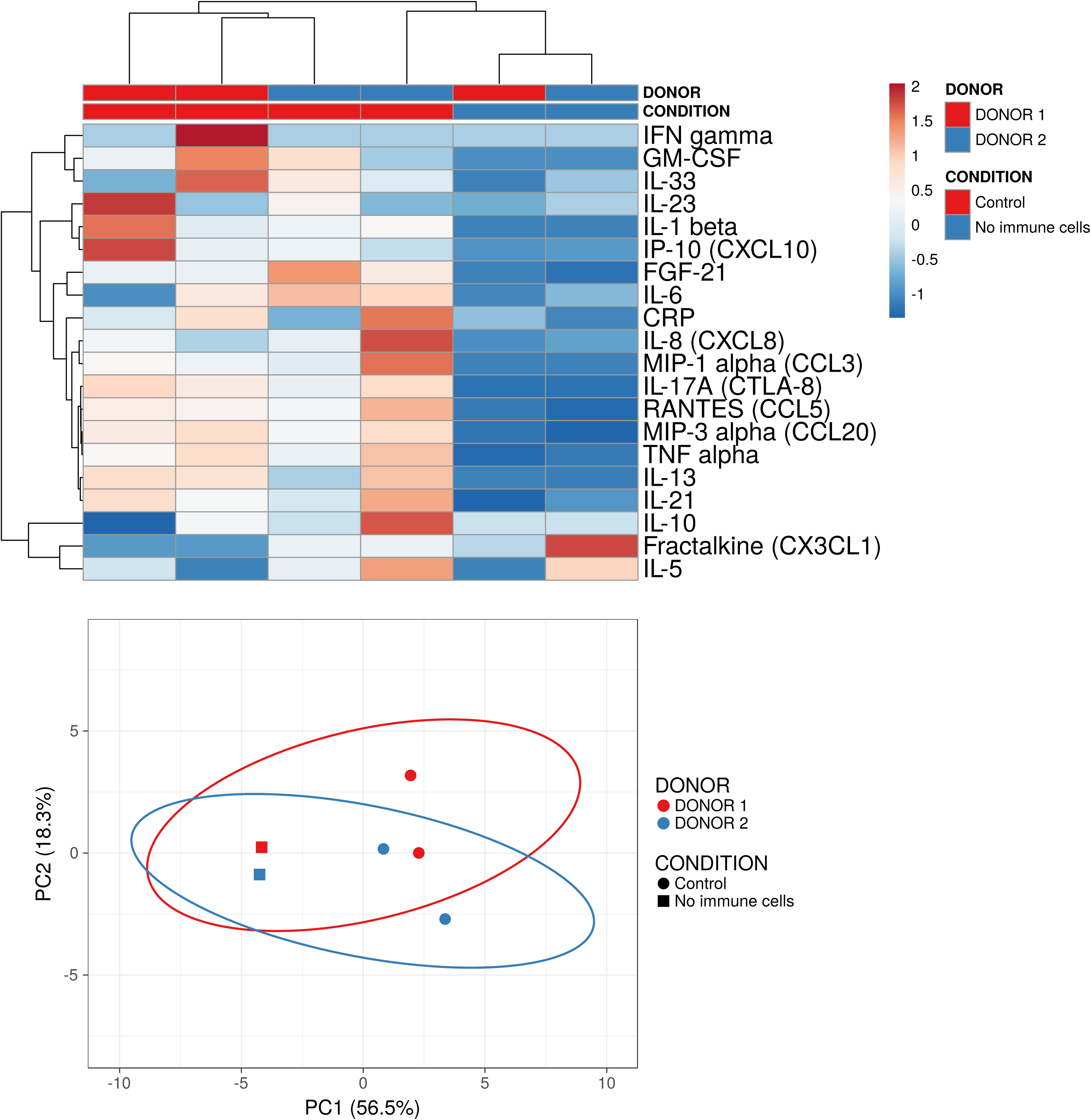
Heatmap and PCA clustering using ClustVis, of measured cytokines and chemokines, during gut-liver interaction studies of parenchymal tissue alone or in the presence of their respective tissue-resident immune cells. Factors were analyzed in shared universal media two days post-interaction. Data represents two separate donors and two biological replicates per donor.

**Figure S4:**
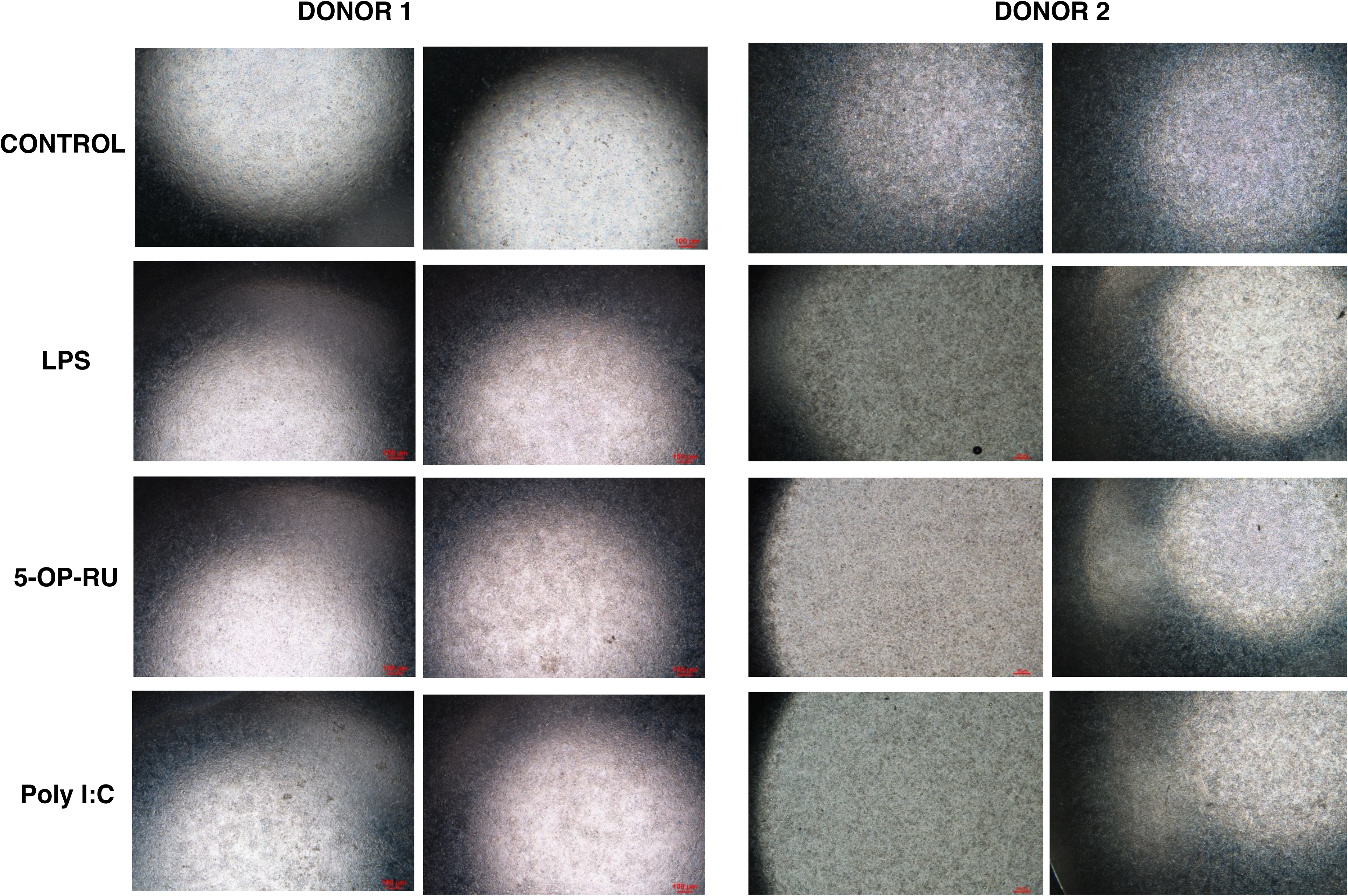
Brightfield images (4x) of intestinal monolayers at day 2 post-interaction and studies of immune activation. Donor 1: first and second columns, Donor 2: thhird and fourth columns.

**Figure S5:**
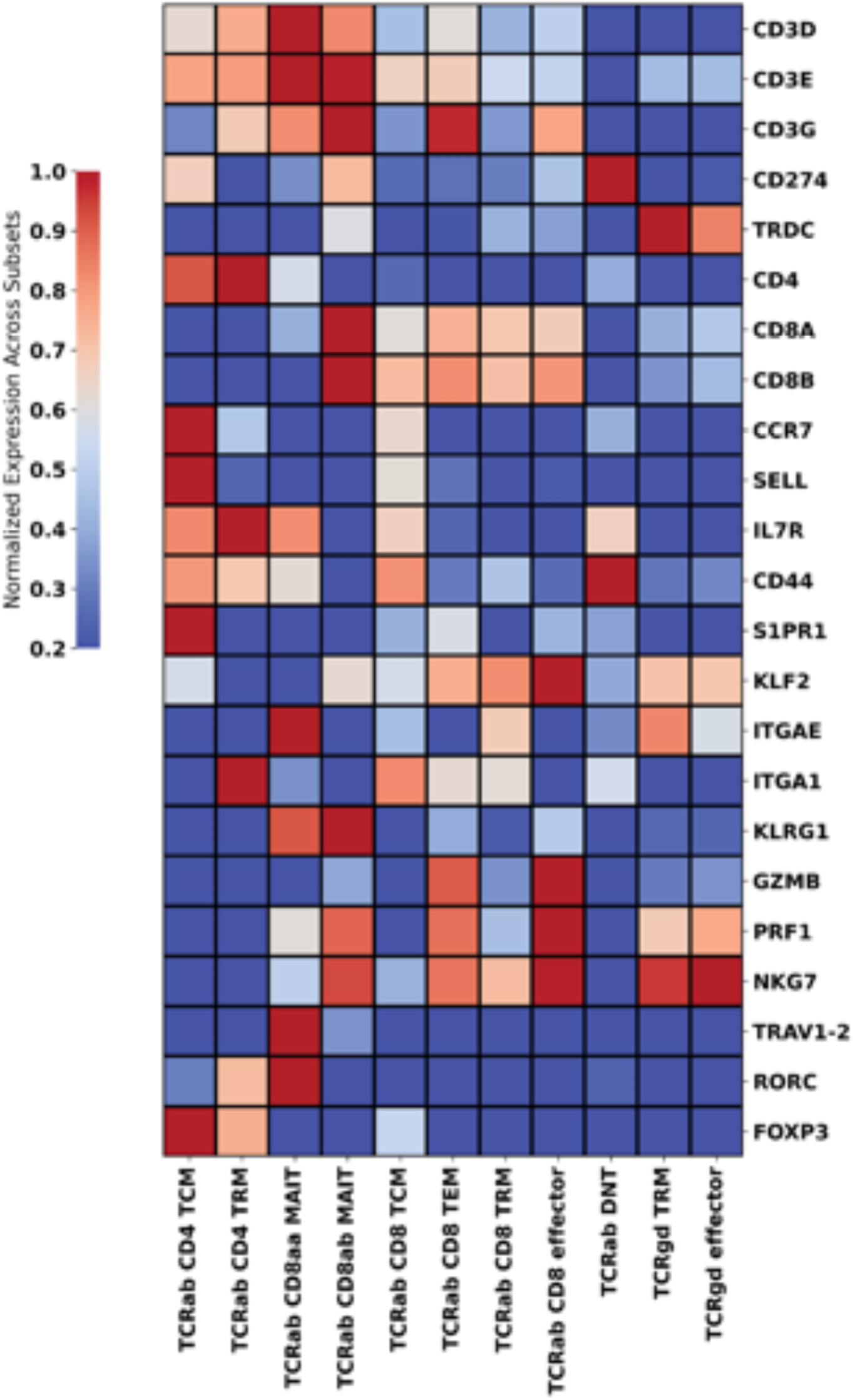
Heatmaps Showing the Gene Expression of T cell subsets markers in all donors’ liver T cells. TCM: Central Memory T cells; TEM: Effector Memory T cells; TRM, Tissue Resident Memory T cells; DN, CD4/CD8 double negative; MAIT: Mucosal-associated invariant T cells.

