## Supplemental figures for "Personalized Gut–Liver Microphysiological System Maps Donor-Specific Tissue-Resident Immunity and Reveals a Conserved Metabolic Crosstalk"

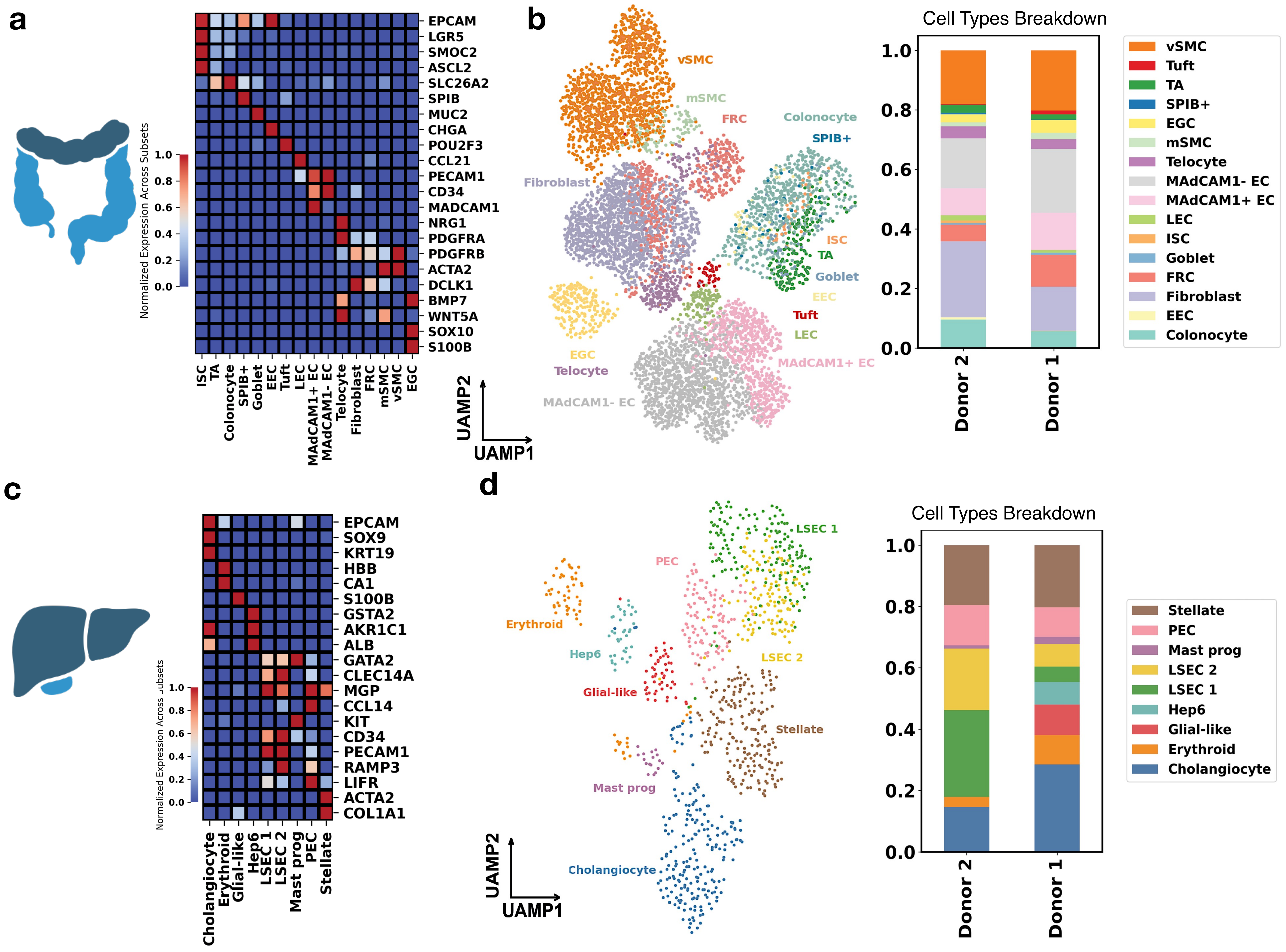


**Figure S1: Single-cell characterization of same-donor colon and liver tissue reveals differences in non-immune cell composition. a,** Marker gene expression in all identified colon non-immune cells. The color of each cell in the heatmap denotes the normalized cell type-averaged expression of each marker. Abbreviations: Colonocyte - colonocytes; EEC - enteroendocrine cells; Fibroblast - fibroblasts; FRC - fibroblastic reticular cells; Goblet - goblet cells; ISC - intestinal stem cells; LEC - lymphatic endothelial cells; MAdCAM1$^+$ EC - MAdCAM1$^+$ endothelial cells; MAdCAM1$^-$ EC - MAdCAM1$^-$ endothelial cells; Telocyte - telocytes; mSMC - muscularis smooth muscle cells; EGC - enteric glial cells; SPIB$^+$ - SPIB$^+$ cells; TA - transit amplifying cells; Tuft - tuft cells; vSMC - vascular smooth muscle cells. **b,** Left: UMAP embeddings of cells collected in the colon of all donors; Right: stacked bar plot shows cell type composition in Donor 1 and Donor 2. **c,** Marker gene expression in all identified liver non-immune cells. The color of each cell in the heatmap denotes the normalized cell type-averaged expression of each marker. Abbrevations: Cholangiocyte - cholangiocytes; Erythroid - erythroid cells; Glial-like - glial-like cells; Hep6 - hepatocytes type 6; LSEC Z1 - liver sinusoidal endothelial cells zone 1; LSEC Z2 - liver sinusoidal endothelial cells zone 2; Mast prog - mast progenitor cells; PEC - portal endothelial cells; Stellate - hepatic stellate cells. **d,** Left: UMAP embeddings of cells collected in the liver of all donors; Right: stacked bar plot shows fractional cell type composition in Donor 1 and Donor 2.

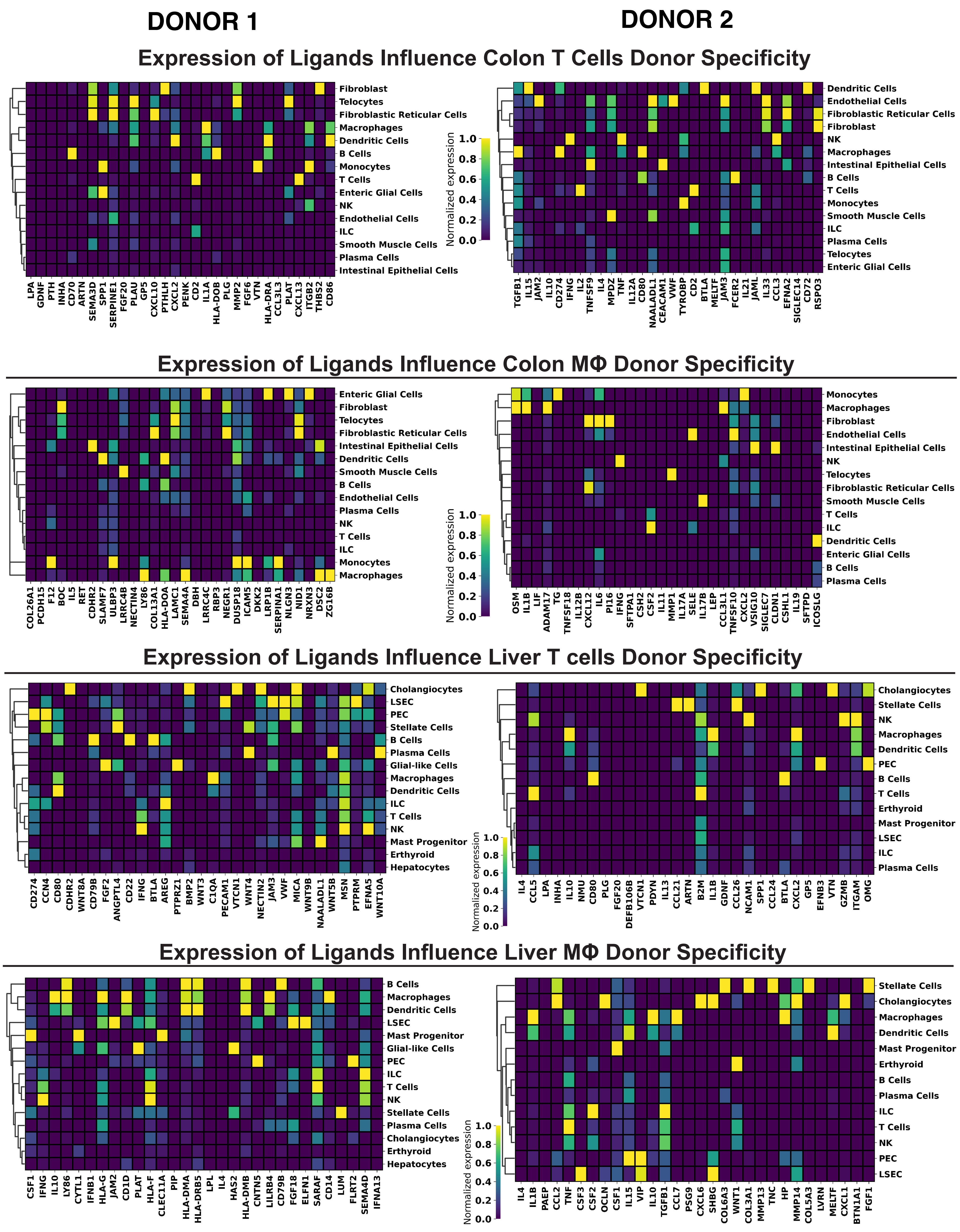


**Figure S2:** Heatmaps showing the Gene Expression of NicheNet-inferred ligands used for AUC Calculation in each donor’s colon and liver cell types.

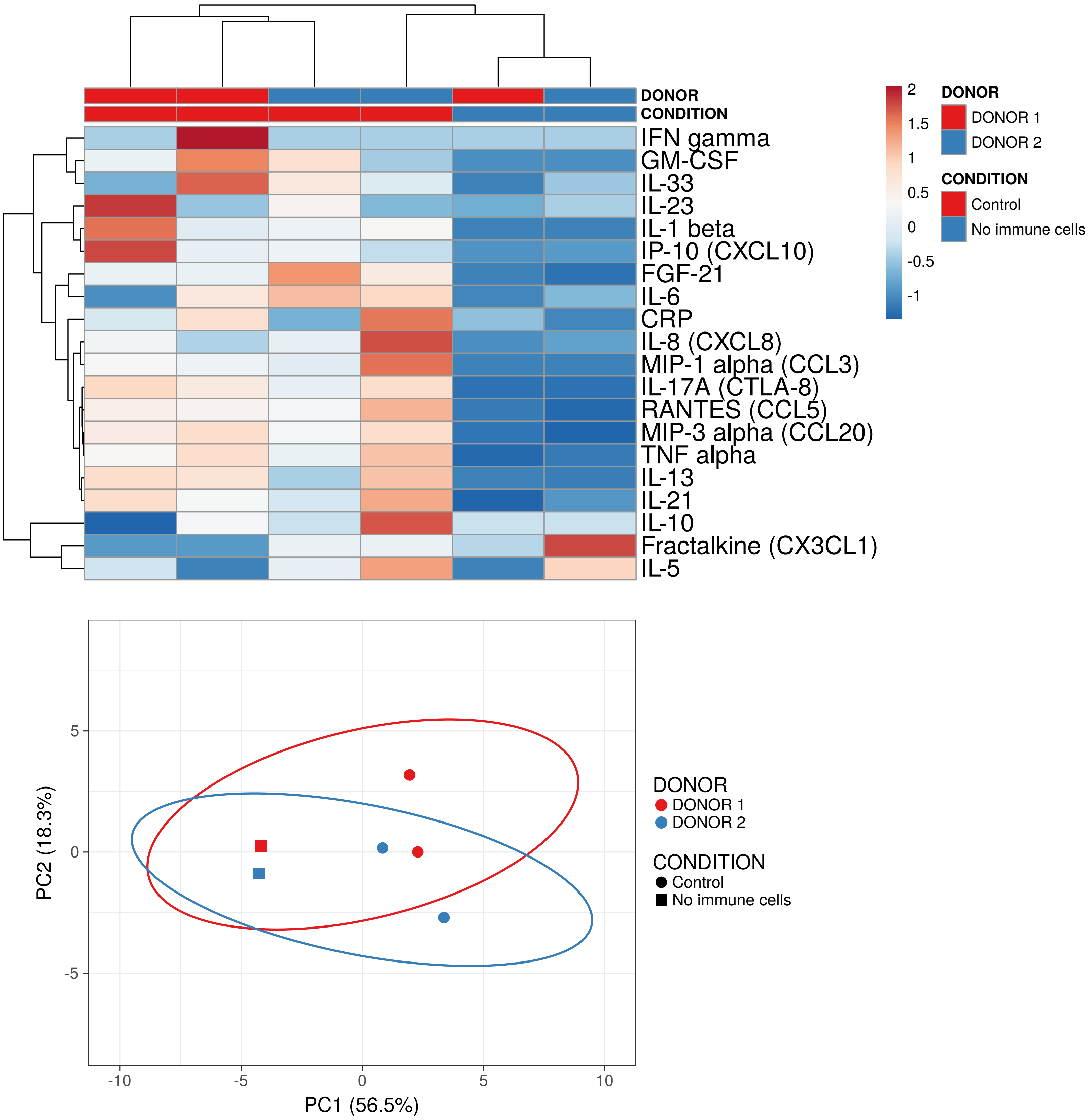


**Figure S3:** Heatmap and PCA clustering using ClustVis, of measured cytokines and chemokines, during gut-liver interaction studies of parenchymal tissue alone or in the presence of their respective tissue-resident immune cells. Factors were analyzed in shared universal media two days post-interaction. Data represents two separate donors and two biological replicates per donor.

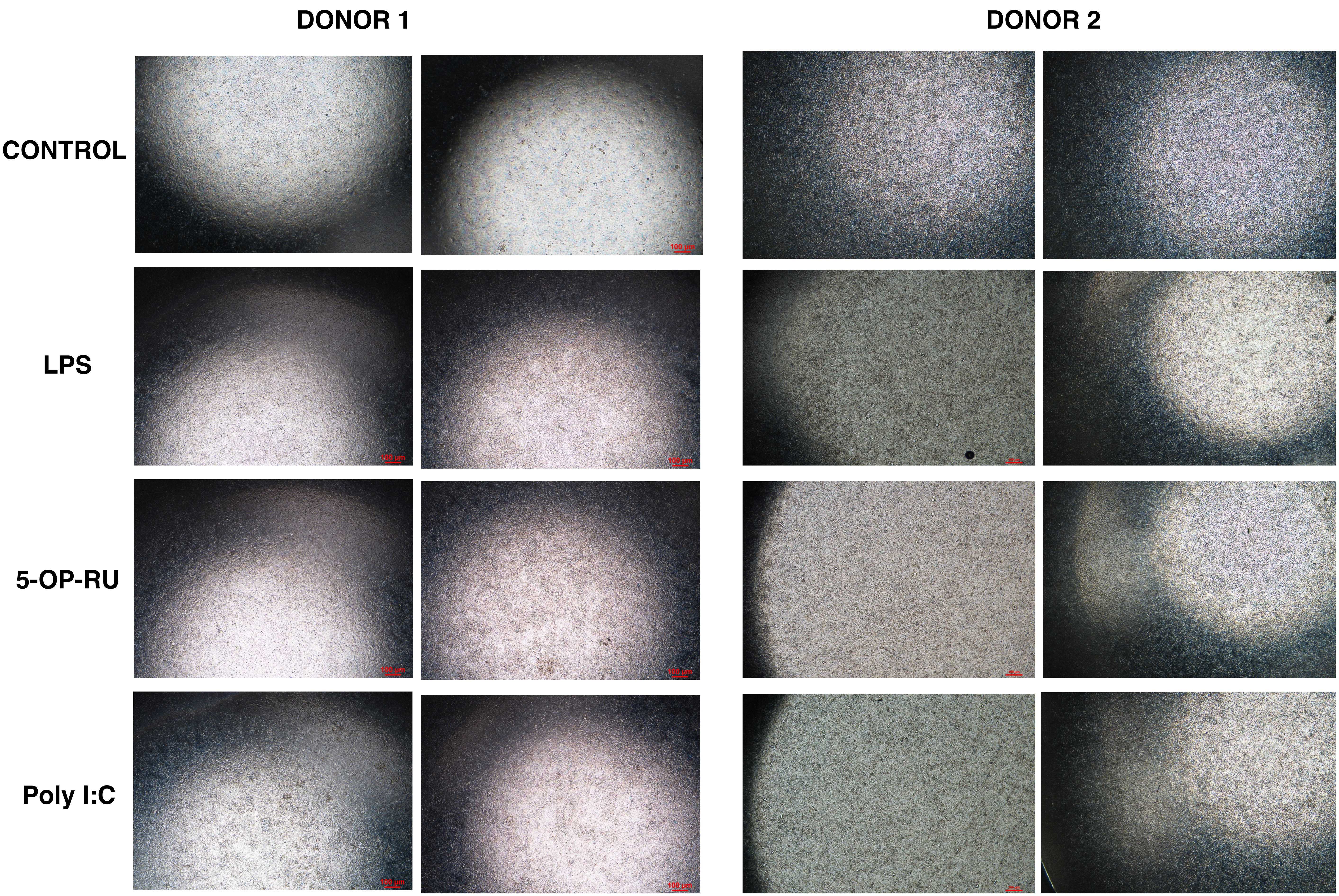


**Figure S4:** Brightfield images (4x) of intestinal monolayers at day 2 post-interaction and studies of immune activation. Donor 1: first and second columns, Donor 2: third and fourth columns.

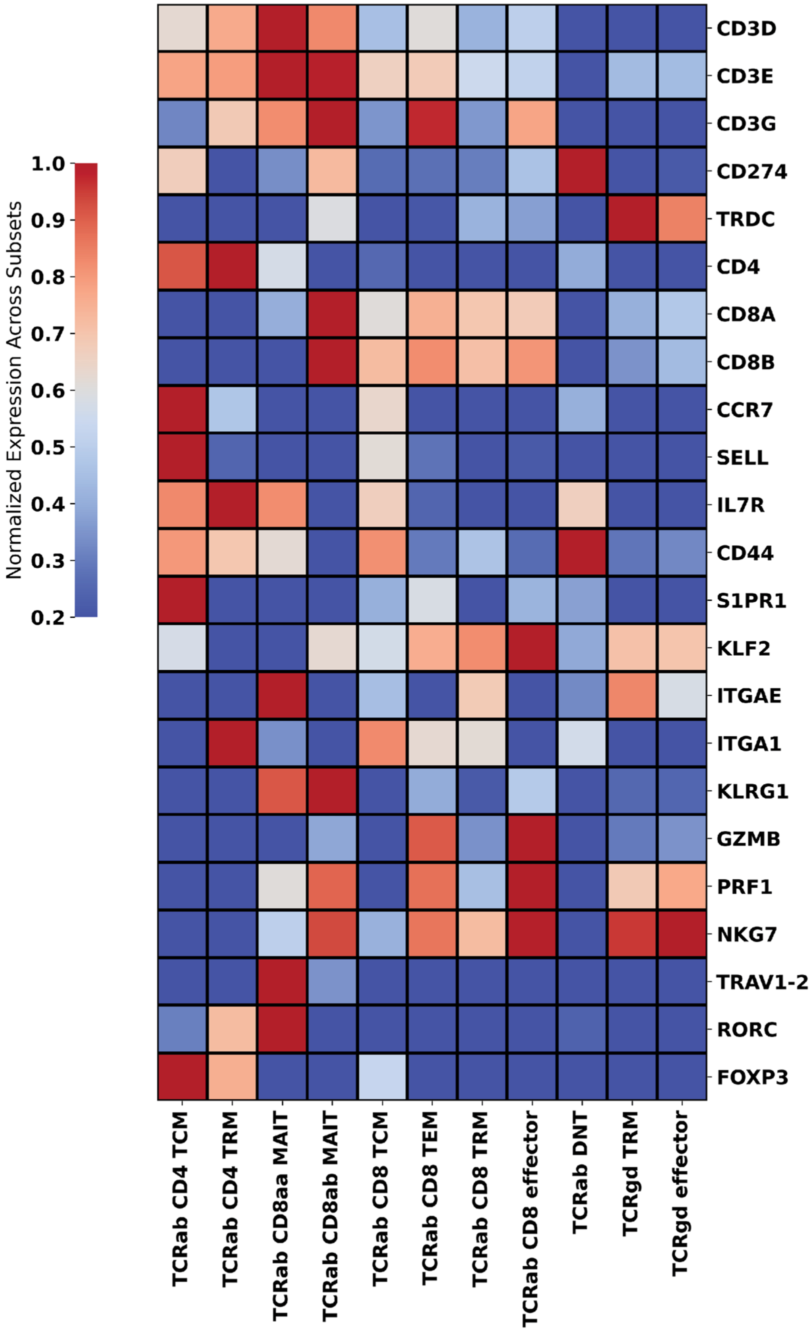


**Figure S5:** Heatmaps Showing the Gene Expression of T cell subsets markers in all donors' liver T cells. TCM: Central Memory T cells; TEM: Effector Memory T cells; TRM, Tissue Resident Memory T cells; DN, CD4/CD8 double negative; MAIT: Mucosal-associated invariant T cells.
